# Design considerations for workflow management systems use in production genomics research and the clinic

**DOI:** 10.1101/2021.04.03.437906

**Authors:** Azza E Ahmed, Joshua M Allen, Tajesvi Bhat, Prakruthi Burra, Christina E Fliege, Steven N Hart, Jacob R Heldenbrand, Matthew E Hudson, Dave Deandre Istanto, Michael T Kalmbach, Gregory D Kapraun, Katherine I Kendig, Matthew Charles Kendzior, Eric W Klee, Nate Mattson, Christian A Ross, Sami M Sharif, Ramshankar Venkatakrishnan, Faisal M Fadlelmola, Liudmila S Mainzer

**Affiliations:** Center for Bioinformatics & Systems Biology, Faculty of Science, University of Khartoum, Khartoum, 11111, Sudan; Department of Electrical & Electronic Engineering, Faculty of Engineering, University of Khartoum, Khartoum, 11111, Sudan; National Center for Supercomputing Applications, University of Illinois at Urbana-Champaign, Urbana, IL 61801, USA; Department of Computer Science, University of Illinois at Urbana-Champaign, Urbana, IL 61801, USA; Department of Molecular and Cellular Biology, University of Illinois at Urbana-Champaign, Urbana, IL 61801, USA; Department of Quantitative Health Sciences, Center for Individualized Medicine, Mayo Clinic, Rochester, MN 55905, USA; Department of Crop Sciences, University of Illinois at Urbana-Champaign, Urbana, IL 61801, USA; Department of Information Technology, Department of Laboratory Medicine and Pathology, Mayo Clinic, Rochester, MN 55905, USA; Laboratory Pathology and Extramural Applications, Department of Laboratory Medicine and Pathology, Mayo Clinic, Rochester, MN 55905, USA; Carl R. Woese Institute for Genomic Biology, University of Illinois at Urbana-Champaign, Urbana, IL 61801, USA

## Abstract

**Background:** The changing landscape of genomics research and clinical practice has created a need for computational pipelines capable of efficiently orchestrating complex analysis stages while handling large volumes of data across heterogeneous computational environments. Workflow Management Systems (WfMSs) are the software components employed to fill this gap.

**Results:** This work provides an approach and systematic evaluation of key features of popular bioinformatics WfMSs in use today: Nextflow, CWL, and WDL and some of their executors, along with Swift/T, a workflow manager commonly used in high-scale physics applications. We employed two use cases: a variant-calling genomic pipeline and a scalability-testing framework, where both were run locally, on an HPC cluster, and in the cloud. This allowed for evaluation of those four WfMSs in terms of language expressiveness, modularity, scalability, robustness, reproducibility, interoperability, ease of development, along with adoption and usage in research labs and healthcare settings. This article is trying to answer, *“which WfMS should be chosen for a given bioinformatics application regardless of analysis type?”*.

**Conclusions:** The choice of a given WfMS is a function of both its intrinsic language and engine features. Within bioinformatics, where analysts are a mix of dry and wet lab scientists, the choice is also governed by collaborations and adoption within large consortia and technical support provided by the WfMS team/community. As the community and its needs continue to evolve along with computational infrastructure, WfMSs will also evolve, especially those with permissive licenses that allow commercial use. In much the same way as the dataflow paradigm and containerization are now well understood to be very useful in bioinformatics applications, we will continue to see innovations of tools and utilities for other purposes, like big data technologies, interoperability, and provenance.

## Introduction

Today’s era of data intensive science is introducing drastic changes to the scientific method^1,2^. Genomics has turned into a large-scale data generation science on par with astronomy and physics^3,4^. With this comes a shift in computational environments from local High Performance Computing (HPC) facilities, to distributed grids, and more recently cloud resources, especially within large-scale multi-center collaborative projects^5^. Likewise, the pressure to process the ever-increasing amount of data at an ever-increasing pace is driving the evolution of software to automate and parallelize analyses in these HPC environments^6^.

*Scientific* Workflow Management Systems (WfMSs) automate computational analyses by stringing together individual data processing tasks into cohesive pipelines^7,8^. They abstract away the issues of orchestrating data movement and processing, managing task dependencies, and allocating resources within the compute infrastructure^9^. Additionally, some WfMSs provide mechanisms to track data provenance, execution errors, user authentication, and data security (Fig 1). The rise of WfMSs in modern science has prompted the creation of new standards in the form of Findable, Accessible, Interoperable and Reproducible (FAIR) principles for tools, workflows, and dataset sharing protocols^10^. These criteria now drive the evolution of containerized software^11,12^ and standard Application Programming Interface (API)s for defining, sharing, and executing code across a range of computational environments^13^.

**Figure 1.**
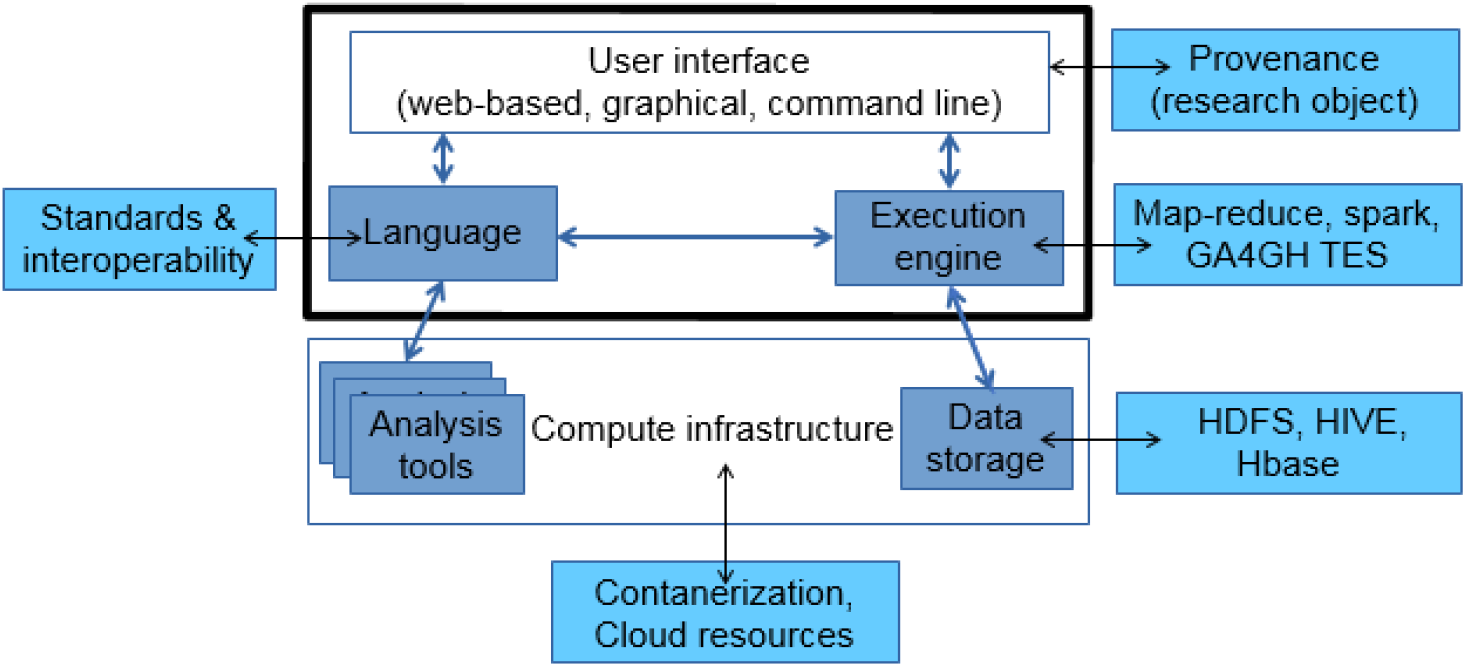
A WfMS is middleware between the analyst and the computational environment. It encompasses the workflow language specifications to interconnect the analysis executables, and the execution engine to dispatch tasks and manage dependencies on the compute infrastructure.

Given that so many groups are implementing and using WfMSs^14,15^, we present a systematic, quantitative evaluation and comparison of their capabilities, focusing on deployment and management of analyses that require complex workflow architecture involving loops, conditional execution, and nestedness. Unlike prior reviews (e.g.,^16^), we will focus on the management of very large analyses across dozens to hundreds of nodes, under circumstances where human interaction would be a significant interruption. These kinds of analyses are performed in large sequencing facilities, major research hospitals, and the agricultural sector.

We identified the following aspects of WfMSs relevant to bioinformatics: (1) *modularity* of the pipeline to enable checkpointing; (2) *scalability* with respect to the number of tasks in the pipeline and the number of nodes utilized per run; (3) *robustness* against failures due to data issues, resource unavailability, or aborted execution; (4) *reproducibility* via logs recording data provenance and task execution; (5) *portability* across compute environments; (6) *interoperability* of metadata and representation enabling workflow registration in common repositories, language standardization, and ability to translate the same workflow into several programming languages; and (7) *ease of development* by users with a range of experience and computational knowledge. We evaluate these aspects both for the purposes of research analyses and their use in clinical settings, requiring data privacy, governance and strict validation of correctness. The results drive our recommendations for using different WfMSs in those settings, and ideas for the future of workflows in biological computing.

## Results

### Philosophy and main purpose of the chosen WfMSs

The usability, features and performance of a WfMS are driven by the purpose for which it has been developed. The Common Workflow Languagen (CWL) is a language specification designed by the bioinformatics community to unify the style, principles and standards of coding pipelines, in a way that is agnostic of the hardware. It prioritizes reproducibility and portability of workflows and hence requires explicit/pedantic parameters definitions, making it very verbose. In contrast, Workflow Description Language (WDL) is a language specification that emphasizes human readability of the code and an easy learning curve, at the cost of being restrictive in its expressiveness (fig Supplementary 4). Nextflow is a complete system that combines the workflow language and execution engine, and is perhaps one of the most mature WfMSs to-date. Desirable features, such as readability, compactness, portability and provenance tracking are available, yet coding is very straightforward, even for a relative beginner in biological computing. Similarly, Swift/T is a complete system: `Swift`, the parallel scripting language, is powered by `turbine`, the execution engine. It was written by physicists and engineers to emphasize scalable deployment of short, rapid-fire tasks at exascale. It is thus a fairly low-level language (similar to C) and extremely powerful, but has a steep learning curve^17^. Below we explore the impact of these different philosophies on the practicalities of using the four WfMSs in production bioinformatics.

### Language expressiveness

Workflow languages are Domain Specific Languages (DSLs) designed to express the architectures of workflows. A *complete* DSL provides the ability to express any workflow pattern^18^ via a rich library of functions, or the means to write custom functions. These abilities, as well as the look and feel of a workflow language, is a function of its parent language (Table 1). Swift/T inherits the flexibility and versatility of C, incorporating all the familiar functions and ability to write custom functions, and drawing from a wide array of pre-existing libraries, including those written in `Tcl`, the parent language of `turbine`. Swift/T reads like a low-level language, which can be difficult for a novice programmer, but provides unparalleled ability to express any complicated workflow logic and embed any advanced algorithm or operation. The Groovy-based Nextflow is similarly powerful, though easier to work with, providing the object-oriented look and feel of Java and access to any library written for JVM. Example implementations of complex patterns, such as upstream process synchronization, exclusive choice among downstream processes, and feedback loops, are available in the documentation^19^. Like Swift/T, Nextflow treats functions as *first class objects*^20^ that can be used in the same ways as variables, enabling the programmer to create easily extensible pipelines, which is a very important feature in the world of ever-changing bioinformatics analyses.

**Table 1.** Summary of language-level differences among Swift/T, Nextflow, CWL and WDL.

| Aspect | Swift/T | Nextflow | CWL | WDL |
| --- | --- | --- | --- | --- |
| Parent language | C, tcl | Ruby and Groovy | N/A | N/A |
| Compilation | Compiled | Interpreted | Compiled | Compiled |
| GUIs | - | NextflowWorkbench<sup>27</sup>,<br>DolphinNext<sup>28</sup> | Rabix composer | Pipeline Builder<sup>29</sup> |
| DSL features | complete, extensible in tcl | complete, extensible in<br>Groovy and Java | limited standard library, extensible via<br>javascript | limited standard library |
| Variables | typed, unique within<br>scope | qualified, unique within<br>scope | typed, unique identifiers | typed, fully qualified names |
| Loops | sequential `for` and<br>parallel `foreach` | parallel queue channels | parallel scatter via<br>ScatterFeatureRequirement | parallel scatter |
| Conditionals | `if-else` and `no-fall`<br>through `switch`<br>statements | via `when` declaration within<br>a process | `when` and `pickValue` fields<br>proposed in CWLv1.2 | `if` blocks producing<br>optional output types |
| Enforcing good<br>practices | - | nf-core<br>(https://nf-co.re/) | CWL guide<br>(https://www.commonwl.org/user_guide/rec-practices/) | - |

CWL and WDL are qualitatively different. They are better viewed as language specifications with strictly defined grammar. Parsers built in other languages, such as Java or Python, interpret this grammar. Thus, CWL and WDL are more restrictive in their expressiveness, but more readable and easier to use. CWL has no functions, but supports Javascript code blocks to express complex code patterns, provided the `InlineJavascriptRequirement` is specified in the script document. However, the CWL team does not consider these code blocks a good coding practice and advises against overusing them^21–23^. Worse yet, conditionals were not directly supported in CWL until version 1.2.0 released in August 2020, after much discussion in the community^24^. Likewise, WDL does not permit programmers to define custom functions and has a very limited library of basic operations. Furthermore, both languages evolve independently of execution engines, which sometimes fail to provide support for certain features. For example, until March 2020, nesting conditionals within loops was not supported with `toil-wdl-runner`^25^, nor are the nested loops in WDL `draft-2` code executable by `Cromwell`^26^ (see table 2), even though WDL specification does not forbid these patterns. Counterintuitively, this makes CWL and WDL particularly well suited for describing biological analysis workflows, by focusing on declarative syntax where each step of the workflow appears clearly in the script. Their expressiveness limitations have been purposefully imposed to enforce good coding practices and prevent unnecessarily complex workflows that cannot be unambiguously resolved. Despite these advantages, experienced coders may find CWL and WDL somewhat claustrophobic.

**Table 2.**
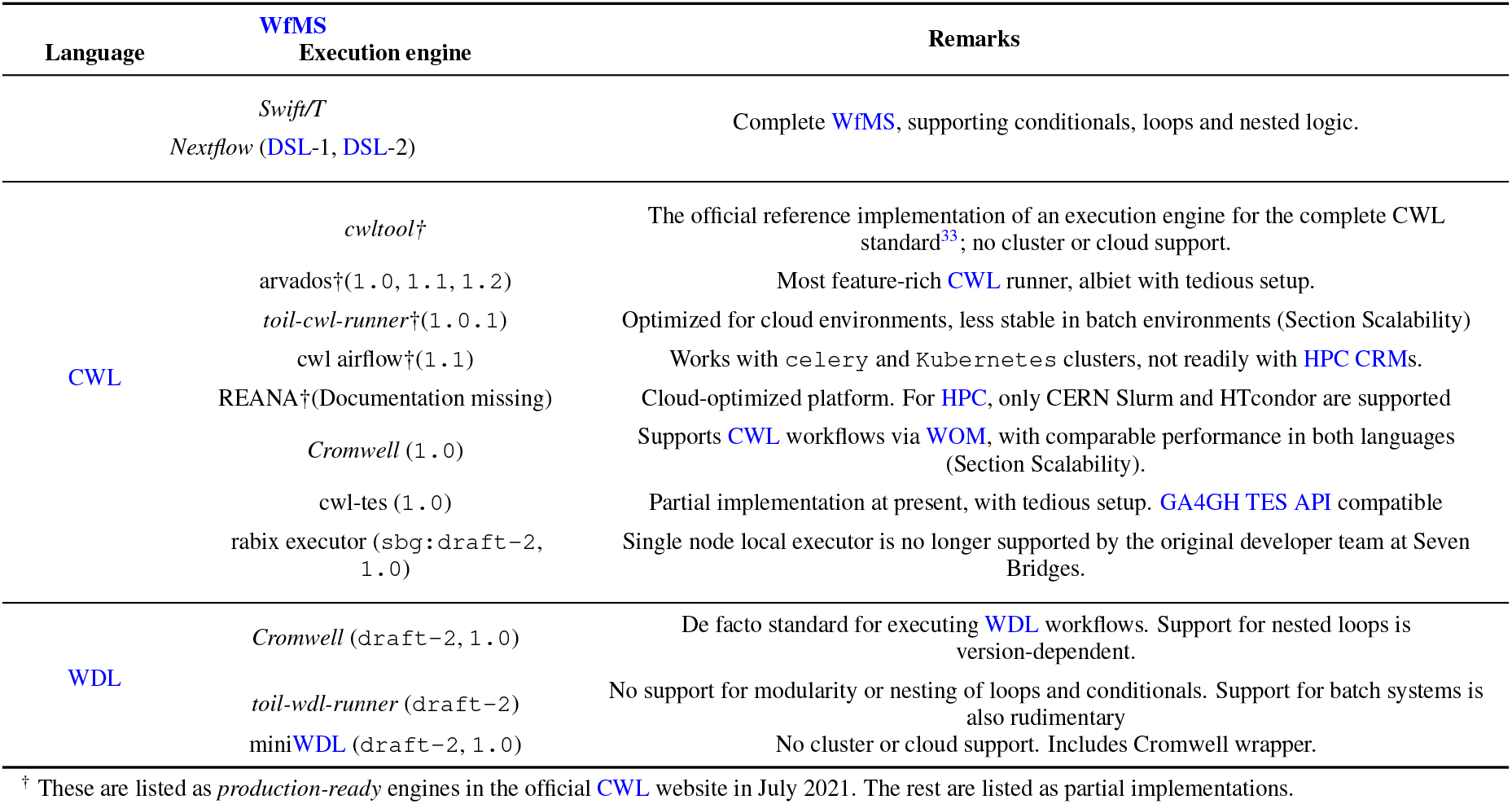
Summary of executor-level differences among Swift/T, Nextflow, CWL and WDL. Any given feature of a workflow language can be assumed supported by the executor, unless we note otherwise. Supported language versions are in parentheses for each executor. *Italics* indicates engines we thoroughly examined.

### Support for modularity

Modularity is a very important design principle for production bioinformatics workflows. The core idea is to build a library of reusable modules (tasks or subworkflows) and assemble them into various master workflows (Fig 2). This enables (1) performing different analyses without having to refactor the entire workflow; (2) check-pointing and restart of a workflow run from a task in the middle of analysis if needed; and (3) customizing runtime environments and compute resources which may vary between analysis stages.

**Figure 2.**
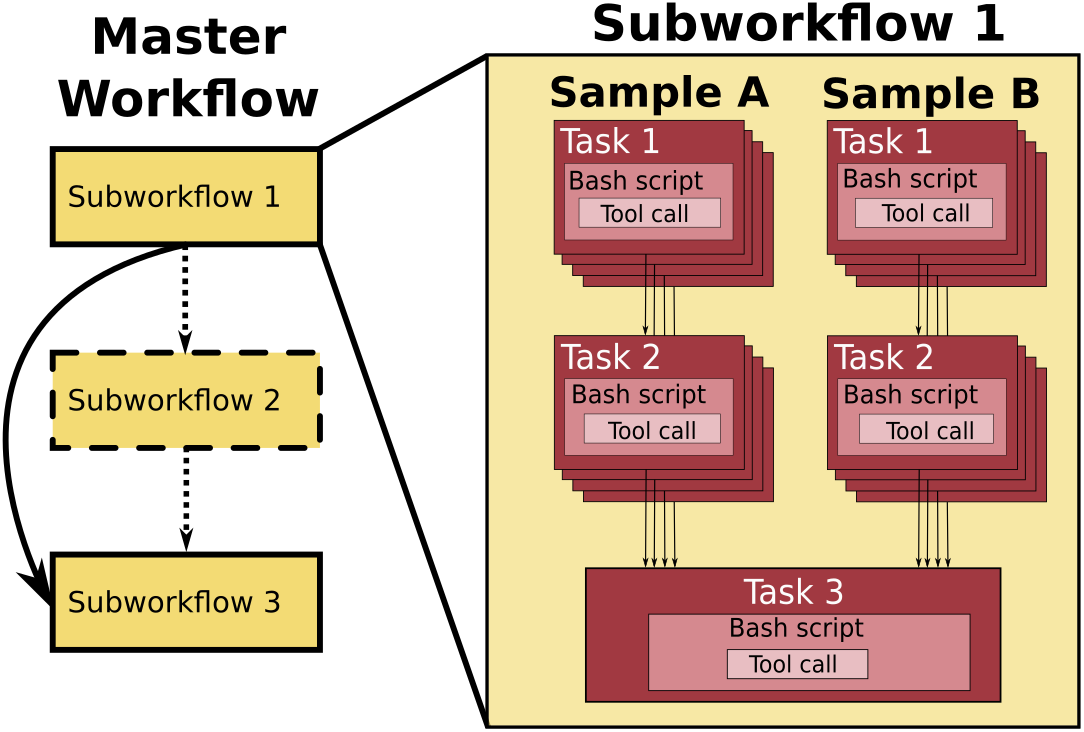
Bioinformatics workflows with multiple levels of complexity warrant a modular construction. It is easiest to program the workflow when its logic is abstracted away (in Tasks, red) from the command line invocations (in Bash scripts, pink) of the bioinformatics tools (light pink). Individual workflows can be further used as subworkflows of a larger Master workflow (e.g., Fig Supplementary 1). This architecture facilitates expression of additional complexity due to optional modules (dashed line), nested levels of parallelism (groups of arrows connecting red rectangles) and scatter-gather patterns (task 2 scattered across samples being merged into task 3).

The superior expressiveness and extensibility of Swift/T make it trivial to implement modularity via user functions, library imports, or `leaf` functions wrapping scripts written in other languages. Out of the four WfMSs we are comparing, Swift/T is the most permissive, at the cost of not having an explicit notion of a workflow. Nextflow is similar, but not quite as permissive as Swift/T, where it defines `process`es to wrap user’s scripts written in other languages and considers a workflow to be a series of those `process` definitions. Nextflow lacked the ability to import and reuse processes, until recently with Nextflow DSL-2. WDL has the most intuitive modular workflow scripting, as it explicitly defines `task`s wraping Bash commands or Python code, and `workflow`s composed of calls to those `task`s. This mechanism makes implementing modular workflows matching to the everyday logic of a bioinformatics analyst. The resultant WDL master workflows contain subworkflows, which consist of tasks, that in turn call Bash code and third-party executables^30^. They are very easy to write and read, and are therefore highly extensible and maintainable. In contrast, this readability aspect cannot be said of CWL. It defines `CommandLineTool` and `Workflow` classes to distinguish individual command-line invocations from the workflow logic calling them. Workflows can be nested by treating subworkflows similarly to `CommandLineTool`s, so long as the `SubworkflowFeatureRequirement: {}` is added into the header. Thus, in principle, a CWL workflow is extensible and modular by the design of the language. Unfortunately, the code ends up being extremely verbose (Fig Supplementary 4), and takes a lot longer to develop than the other three WfMSs.

We conclude that all four evaluated languages deliver satisfactory support for modularity at the code level, though ease of use remains in the eyes of the programmer. Use of modularity for custom resource allocation, check-pointing and auto-restart from the point of failure, is the executor’s job, and (sections *Job execution: resources provision* and *Robustness*).

### Data dependencies and parallelism

In the dataflow paradigm^31^, data dependency and parallelism go hand in hand. Blocks of code that do not have data dependencies among them are executed in parallel (implicit parallelism), e.g., quality checks on a BAM file. A different mechanism is usually implemented for code blocks meant to run in parallel (explicit parallelism), e.g., read alignment across multiple lanes (Fig Supplementary 1). The four languages studied here differ in handling the switching between the implicitly parallel and the explicitly serial phases of an analysis (Fig Supplementary 4).

Swift/T employs `foreach` and `for` statements for parallel and sequential iteration over array elements, respectively. The `=>` or `wait` statements enforce serial execution of tasks where explicit data dependency is missing, as `Turbine` will otherwise attempt to parallelize such statements. Yet, with its low-level language style, the vigilance in writing complex Swift/T workflows can be taxing, and the resultant code difficult to debug for parallelism issues.

In contrast, WDL and Nextflow stylistically separate the areas where sequential execution of multiple commands is permitted. command blocks inside WDL `tasks` are equivalent to the `script` blocks inside Nextflow `processes`. Parallel execution is assumed among WDL tasks, unless data dependency exists between inputs and outputs of different tasks. Explicit `scatter` statements parallelize execution over array elements, while results are implicitly gathered. Uniquely, Nextflow defines both process dependencies and parallelization via input/output variables, where a multi-valued `queue channel` signals parallelization over its elements. This elegant approach yields compact code, at the expense of readability. First, it requires careful pipeline design, because a process is executed as many times as the size of its shortest `queue channel`, and their types and sizes matter. Second, gathering results after parallelization needs to be coded explicitly. On the plus side, channels make it trivial to expand pipelines. For example, expanding from single sample to multi-sample joint calling is achieved by merely adding the downstream JointGenotyping process, without the addition of a nested loop across the samples (Fig Supplementary 1).

CWL is fundamentally different: its `CommandLineTool` is an invocation of a single shell command, not a series of sequential commands or even a string of piped commands. Because many tools are common among bioinformatics pipelines (e.g., samtools), this restriction encourages reuse of the corresponding `CommandLineTool` modules, facilitating standardization and therefore reproducibility. It is easy to think of a CWL `CommandLineTool` as a very restricted version of a WDL task: they both have inputs, outputs, metadata, resources options and a script, but in CWL only a single command is allowed. Parallelization in CWL is accomplished via `ScatterFeatureRequirement {}`, similar to scatter blocks in WDL.

### Executor-level differences

The workflow executor is the WfMS component resolving the workflow syntax into a graph of dependencies between tasks, typically expressed as a Directed Acyclic Graph (DAG) (Fig 1). Then it deploys those tasks in the correct order on the given infrastructure by scheduling the jobs, provisioning compute resources, and tracking the jobs to completion. Executors may have other functionalities for data staging, monitoring, and error recovery. These aspects are explored in subsequent sections for key executors of each WfMS (Table 2). In this study, we focus on *production-ready* executors that work in HPC settings. While we do examine portability, and comment on cloud-friendliness, a detailed analysis of runners primarily dedicated to those environments is beyond our scope

In Swift/T and Nextflow, the workflow language and its executor are packaged together, and therefore co-evolve without compatibility issues. Conversely, CWL and WDL only specify the language syntax, which may be supported by a variety of execution engines. This results in a healthy competition among the engines, but also raises compatibility issues.

In addition to standalone executors, there are API libraries for interpreting workflow languages. For example, miniWDL^32^ is a local runner for WDL and a Python API - a developer toolkit enabling WDL workflows to run from within Python scripts. This opens up possibilities of building a richer workflow ecosystem through embedded data parsers, job visualization, and other useful features.

### Workflow dependency graph resolution and visualization

The dataflow paradigm requires the executor to deterministically resolve the supported workflow patterns^18^ into unambiguous DAGs while controlling for environmental variables and random seeds^34^. This determinism ensures that all processes using independent inputs are scheduled to run in parallel; whereas processes linked via data dependencies are scheduled to run in the appropriate order. Due to these requirements, some features, e.g., conditionals^24^ and workflow dry runs^35^, are difficult to implement in dataflow programming. Each execution engine we studied negotiates its own semantics with the corresponding workflow language to ensure correct DAG construction, succeeding in its own way. Workflow DAG resolution requires handling many small details and careful development and co-evolution between the executor and the workflow language. Conformance of the executor to the language specification is critical to avoid unresolvable patterns when programming complex workflows or when migrating from one executor to another^36^.

Among the four WfMSs we considered, all but Swift/T have built-in engine functionality or auxiliary tools to visualize DAGs for debugging and documentation. Nextflow produces the DAG upon completion of a workflow execution, in static or interactive format, using Cytoscape.js^37^ and Dagre graph visualization libraries (Fig 4a). Alternatively, NextflowWorkbench^27^ and DolphinNext^28^ provide convenient graphical and web interfaces developed outside the Nextflow core team for visualizing, creating, deploying and executing Nextflow pipelines. For CWL, CWLViewer^38^ conveniently produces the DAG of a workflow script from its GitHub repository (Fig 4e), if the code is public and complies with CWL best practices. The Rabix suite, by Seven Bridges, provides powerful CWL interactive visualization library (`CWL-SVG`), GUI (`Composer`), and language server (`benten`) (Fig 4d), which are used in other sophisticated projects like VueCWL^39^. For WDL, besides the *de facto* `womtool` utility and its `graph` visualization option (Fig 4c), EPAM systems have developed Pipeline Builder^29^, a Javascript library for interactively constructing and visualizing WDL scripts (Fig 4b).

### Job execution: resources provision

Bioinformatics workflows are often heterogeneous in terms of the computational resources required for each task. For example, genome assembly can begin with read alignment (a core-intensive process), followed by a deBruijn graph construction (a RAM-intensive process). For efficiency, the executor must decipher which tasks can be run as individual computational units on hardware (i.e. node) with RAM and cores appropriate to needs. Within the dataflow paradigm, these independent units are, by definition, the vertices on the DAG. Upstream and downstream nodes on the DAG (i.e. prerequisites and dependents, respectively) can be placed on other hardware with adequate resources.

The actual provisioning of the computational resources is achieved by interacting with a cluster resource manager (CRM), such as PBS Torque, Open Grid Engine, or Slurm. The CRM tracks the available queues, the available nodes, and how long nodes will remain occupied. If the workflow language has the ability to specify resources necessary for different tasks, then the execution engine may be able to negotiate these requirements with the CRM. This is commonly achieved via executor backends, which provide a mechanism to specify the computational requirements as part of command line workflow invocation or via configuration files.

Particularly useful is the support for *dynamic job scheduling*, present in all four WfMSs under consideration. During workflow execution a job may pass runtime parameters to another job or schedule other jobs. Customization of runtime parameters per workflow stage (section *Nomenclature*), including docker images specifications, memory, queue and/or cloud resources is readily possible in Nextflow, CWL and WDL. In Swift/T however, these details are specified once at the beginning of the workflow; thus, customization can only be achieved by breaking up the main workflow into independent pieces and running those smaller workflows independently. Tables 3 and 4 in Ahmed *et al*.^17^ show backends supported by select executors, and Fig Supplementary 3 shows typical command line invocations.

**Table 3.** GitHub activities from each WfMS (March 4th, 2021). *Contributors* is the number of contributors in each repo, *Open and Closed* refer to the count of open and closed issues and pull requests in the repo.

| WfMS | First commit | Contributors | Closed | Open | License |
| --- | --- | --- | --- | --- | --- |
| Swift-t | 2011-05-11 | 16 | 109 | 81 | apache-2.0 |
| Nextflow | 2013-03-22 | 81 | 1770 | 159 | apache-2.0 |
| CWL | 2014-09-25 | 62 | 667 | 249 | apache-2.0 |
| WDL | 2012-08-01 | 44 | 376 | 50 | bsd-3-clause |

**Table 4.**
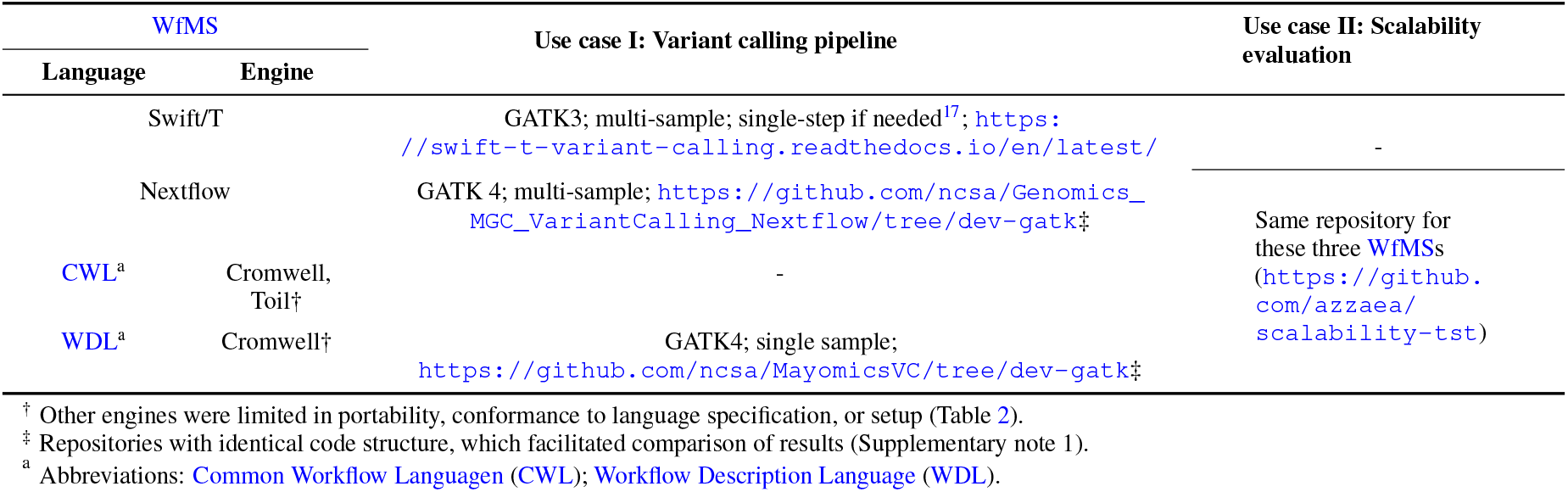
The WfMSs examined in this study.

### Job execution: data staging

Bioinformatics data processing frequently involves movement of private and very large datasets (TBs or more) across infrastructure that is set up on a shared filesystem. The resulting security and performance concerns create a need to isolate the workflow execution environment and provide a means for checking the integrity and permission settings on files used and produced within a workflow run. Data isolation is commonly accomplished by enabling special treatment of the `file` type variable by the engine, e.g., file integrity checking and hashing. Additionally, localization (staging) of inputs into a working directory unique to each computational unit (1) assures that raw input data remain intact; (2) prevents race conditions if a file needs to be accessed by multiple computational units simultaneously; (3) serves as a record of provenance for each input, output, intermediate file, script and log, enabling easy monitoring and debugging; and (4) enables workflow restart from the failed stage without repeating prior computations.

Nextflow and executors of CWL and WDL all provide this staging capability via a canonical hierarchy of execution folders. The working directories of subworkflows, tasks and `Scatter`ed blocks are nested within a parent directory of the run. Unique folder naming is ensured by using long hexadecimals, names of the workflow stages, and/or execution timestamps. The exact structure of the working directory and the subfolder names within vary by engine (Supplementary note 4), and the user has no control over these parameters, which may seem constraining. However, it pays off in permitting the engine to automatically follow the task dependency string and prevent filename clashes for subsequent tasks. Conversely, in Swift/T, the programmer must manually create a directory tree and name files, which is error prone in complex workflows.

In addition to the enforced separation of files in the output folder tree, further data staging can be achieved by placing them on different filesystems, such as a cloud bucket for inputs vs. local folder for outputs. Nextflow supports this out of the box: the analyst need only specify a URL to enable reading of inputs from AWS S3 storage or Google cloud buckets. Cromwell and Toil similarly support this ability, though in Cromwell the programmer needs to be specific about the filesystem being pointed to. Unfortunately, in Swift/T the support is more limited: documentation stipulates the means of specifying remote filesystems, but our experiments with that have not been successful.

### Portability across HPC environments and the cloud

Modern biomedical research increasingly benefits from multi-site collaborations. Support for portability, the ability to run a pipeline in computing environments besides the one on which it was developed, has become one of the deciding factors in adopting a particular WfMS. One of the most important aspects impacting portability is hardcoding any system-specific parameters or paths. Separating the pipeline code from the input specification helps detect and eliminate this problem, usually via configuration files. Nextflow, CWL, and WDL made this a requirement. CWL and WDL imposed further constraints by using structured `YAML` and/or `JSON` files and enforcing variable checks on identifiers or fully qualified names at compile time. Swift/T is the least restrictive, putting the onus on the programmer to ensure that variables are defined in a way that does not impede portability (Fig Supplementary 3).

All executors we examined have ample support for running in a variety of compute environments (Tables 3 and 4 in Ahmed *et al*.^17^), except that support for AWS and Google Cloud Platform (GCP) is insufficient in Swift’s `turbine`. Cloud deployment in general comes with different considerations than HPC. First, the executor needs to communicate with cloud APIs to provision and administer the resources specified in the configurations options. Second, the provisioned cloud resources are typically clean machine instances, providing only the basic operating system and minimal libraries. Thus, executors rely on containerization of software used by the workflow and expect container images of those tools and their dependencies as part of the workflow runtime options. Third, significant cost savings can be achieved when executors support automatic sizing of cloud resources, enabling instances to be spun up when needed and shut down when idle; AWS batch is a good example. Finally, workflows intended to run in the cloud also expect data to be stored in the cloud. Thus, some level of security is expected from the cloud provider. These are important considerations when evaluating the capabilities of a WfMS for a project intended to run in the cloud.

While executors in the cloud may be affected by a variety of other factors, such as provider, zone, availability of resources or type of machine instance, this is an actively developing area of technology and providing a detailed review is beyond the scope of this work. We did, nonetheless, perform a cursory evaluation of portability by deploying the variant calling pipeline in both AWS `Batch` and `Cloud cluster` (Table 5), which was successful.

**Table 5.** Computational testing environments. I and II refer to: *Use case I: variant calling pipeline* and *Use case II: scalability evaluation*.

| WfMS |  | Biocluster | AWS |  |  |
| --- | --- | --- | --- | --- | --- |
| Language | Engine |  | Batch mode | Cloud cluster | Parallel Cluster |
|  | Swift/T | I, II | - | - | - |
|  | Nextflow | I, II | - | I | II |
| CWL | Cromwell | II | - | - | - |
|  | Toil | II | - | - | - |
| WDL | Cromwell | I, II | I | - | II |

### Scalability

When running a workflow on a substantial number of samples in a shared environment, one must weigh the trade-offs between total run time and computational cost, which is a product of both the run time and number of nodes utilized. Two performance metrics are important: (1) How well the execution engine scales with the number of parallel tasks: The task management overhead may increase out of proportion with the total number of tasks, if the engine is not programmed efficiently. (2) How well the engine packs tasks on a node: It must weigh the core and RAM availability on a node against requirements of the tasks, and pack tasks optimally onto nodes, without many gaps or unused resources.

We compared the performance of Nextflow, toil (running CWL), and Cromwell (running WDL and CWL) against these criteria by designing very simple one-step and two-step workflows (Fig 4). A task was just to echo the hostname (i.e. node ID) of the node where the task was placed during execution. The command requires a negligible amount of RAM, only 1 core, and takes a minuscule amount of time. Thus any overhead on task management is readily apparent from the total time of the workflow. We ran the same workflow with varying number of tasks meant to be computed in parallel, on either: 1) the shared 5 nodes HPC, Biocluster, where the effects of queuing can be noted (72 cores each); or 2) a dedicated, fixed-size AWS Slurm `Parallel cluster` of 100 nodes (96 cores each), in an attempt to control for any extraneous performance variation.

On AWS, Nextflow and Cromwell+WDL both showed excellent performance up to about 100 tasks: the elapsed workflow run time did not increase significantly with the number of tasks, suggesting that they were properly parallelized with minimal management overhead (Fig 3, left panel). Nextflow performed particularly well, finishing the runs about 4 times faster than Cromwell. Two-step workflows predictably ran longer, but not twice as long, which indicates a substantial amount of time is spent at startup for both engines. Performance began to break down at higher task counts, intermittently resulting in failure to start jobs. Nextflow could not be used at all on more than 512 tasks: it quickly stopped the run and cleaned-up. In contrast, Cromwell became unusable beyond 1024 tasks, but the clean-up for the failed job took a very long time (hours in some cases) before finally reporting an exit code.

**Figure 3.**
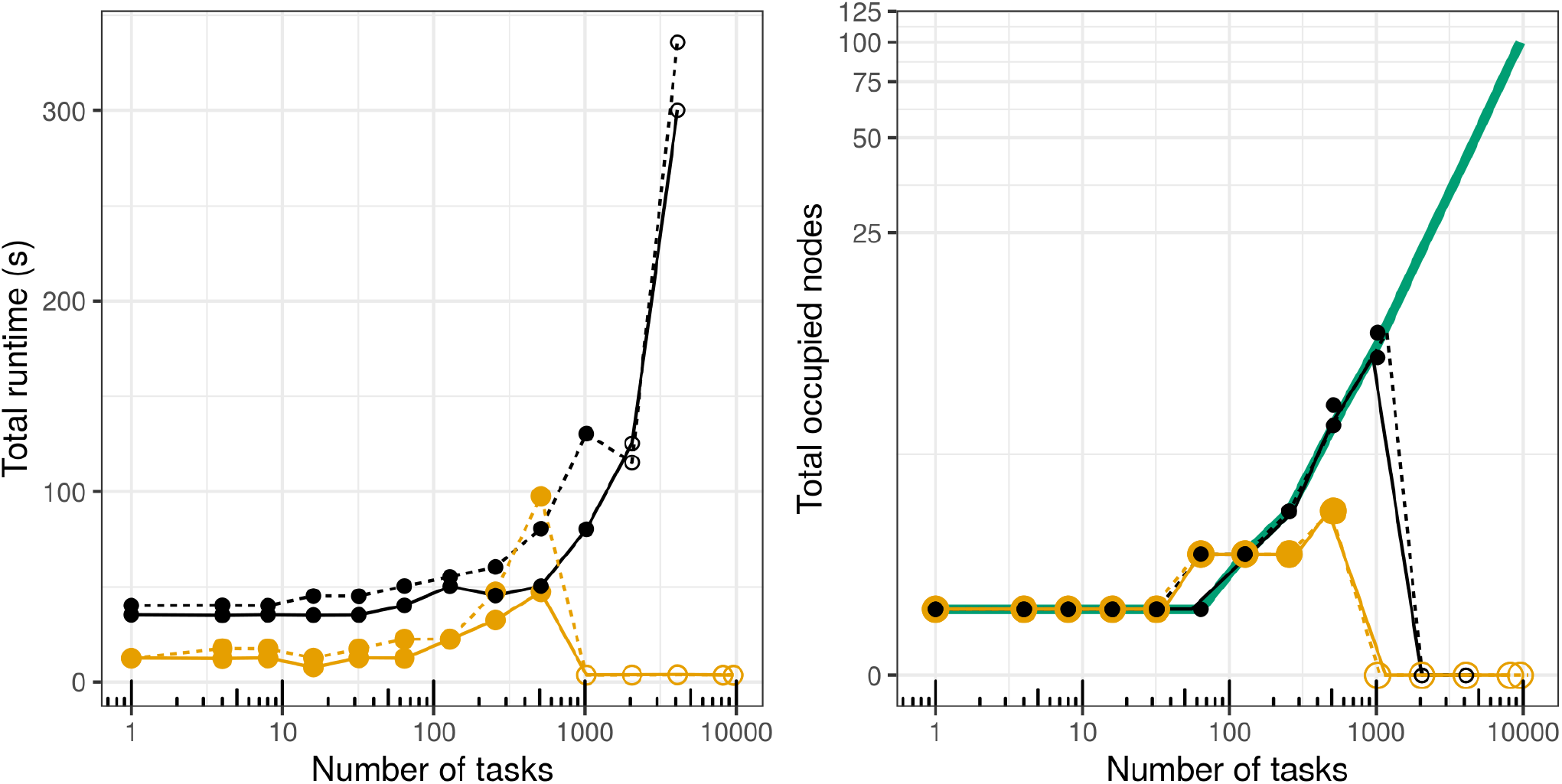
Scaling a one-step (solid line) and two-step (dashed line) workflow in Cromwell+WDL (black) and Nextflow (yellow) on AWS `Parallel cluster`. The thick green line in the right panel is the theoretical optimum of the number of nodes to be occupied by the tasks, computed as the ceiling of tasks/cores-per-node (96). Empty circles denote failed runs.

In subsequent experiments on Biocluster, repeated 5 times to account for queue variations, we used more recent versions of both engines, in addition to Cromwell+CWL and toil+CWL (Supplementary note 5.2). We excluded Toil+WDL as it did not readily support Slurm CRM. Again, Nextflow always ran much faster than the others, up to 50x at times. Toil+CWL, however, failed rather randomly and inexplicably, but quickly, at different scales in each run. Cromwell+CWL performed similar to Cromwell+WDL, with no failures in the tested range.

To understand what caused the scaling issues, we looked into process context switches, both voluntary (where processes yield CPU access to another) and involuntary (where the kernel suspends process access) and found both types to increase with the task count (Fig Supplementary 5, Supplementary 8). CPU utilization was measured as well, as the `user+system` time divided by the total run time of the task (from the `Linux` time command), however no easily interpretable pattern emerged (Fig Supplementary 6, Supplementary 9). Notably though, Cromwell tends to have better CPU utilization with increased task count, while toil+CWL is more stable across the range. Additionally, with the more recent versions of these runners, speed gains and better CPU utilization metrics were realized (Supplementary notes 5.2 and 5.3).

We measured the quality of node packing by using the outputs from our mini workflows, the hostnames where tasks were run. These records were deduplicated, giving the count of individual nodes used by the workflow. We expected no re-use of nodes when the task count is less than the total core count (as all tasks are immediately parallelizable in this case); but queuing of tasks otherwise. The theoretical expectation of the used number of nodes is marked as the thick green line in Fig 3 and Fig Supplementary 7 (rightmost panels), where values below the green line suggest queuing of processes; while values above it suggest the executor is using more nodes than necessary.

Indeed, on AWS, the two engines correctly placed all tasks onto the same 96-core node in runs up to 32 tasks (Fig 3, right panel). In this experiment, while we were controlling Nextflow’s `maxForks` directive, we used the default value of 100 for the `queueSize` parameter, which defines the number of tasks it handles in parallel. This resulted in Nextlow unnecessarily constraining tasks ≥ 256 to to less than 3 nodes, as would have been optimal. Yet, this inefficiency did not prevent Nextflow from outperforming Cromwell in terms of run time, despite Cromwell spreading the processes across nodes in near-perfect alliance with theoretical expectation. On Biocluster, we controlled both directives in tandem, and this constriction is only observed when the task count (512) exceeded the cluster’s total core count (360) as expected (Fig Supplementary 7, righmost panel). In both experiments, we note patterns of unnecessary spread of tasks among nodes with both engines at times. This is something to keep in mind when working with large data batches, as it is desirable to minimize data movement between nodes^40,41^. A helpful directive to this effect in Nextflow is `scratch`, and in Cromwell is `localization_optional`.

### Robustness

When running large scale analyses, especially in medical production settings, where a workflow failure can lead to delay in diagnosis and treatment of patients, it is extremely important to have a WfMS that facilitates the development of easy to debug and maintain error-free code, and which results in robust execution against variations in the nature of the data, load on the compute system, and hardware failures. Beyond traditional provisions for code robustness, WfMSs have the potential to facilitate recovery and restart after failure of an individual analysis step. Ideally, the need to rerun costly analyses from the very beginning could be obviated via *“safe crash”*: by moving the completely processed files to their destination, deleting partially processed files, and saving execution logs and status (check-pointing) for all parallel processes. A number of approaches have been developed in this field to facilitate these traditional and non-traditional robustness aspects.

*Variable typing* facilitates earlier discovery of bugs before the workflow is run, especially in *compiled* languages (Table 1). While Swift/T, CWL, and WDL provide the typical `String`, `Integer`, `Float`, and `Boolean` types, Nextflow does not distinguish between these but rather uses *qualifiers* to indicate how variables are to be handled. For example, variables local to a process have a `val` qualifier, whereas environment variables should be declared as `env`. Workflows in bioinformatics usually operate on files, thus WfMSs must also define a `file` variable, ideally with a mechanism to check whether the file exists and has the right permissions, to avoid data access failures in the middle of analysis (cf. section *Job execution: data staging*). While the four WfMSs studied provide this, Nextflow goes above and beyond generic functions for reading and writing. Nextflow provides refinements to handle especially large, binary and compressed files. Additional domain functions include counting the number of records in FASTQ/FASTA files, splitting file entries based on chunk size, memory limit, etc. Such well-vetted, built-in functionality significantly reduces the likelihood of programmer error, thus conferring robustness.

*Provisions to ease parsing and validating* the code can greatly contribute to the robustness of the final software. Unsurprisingly, most WfMSs make use of such tools. Nextflow workflows can use `nf-core schema` commands^42^. Similarly, `WOMtool, miniWDL` and `Oliver` ease validating, parsing and generating WDL scripts and inputs. CWL takes advantage of standard editor plugins for vim, emacs, VScode and atom, and code generators for R, Go, Scala, and Python. Swift/T can only be accessed via the command line. No helper library is mentioned in the documentation either.

*Data streaming* is an alternative to data staging, and is very popular in bioinformatics. Here, instead of saving output of an upstream task to a file that is read by the downstream task, data are streamed, usually via a Linux pipe, from one process to the next. Such streaming requires synchronization of the two processes and could lead to complicated logic. Additionally, it can make it harder to record and debug execution logs. Perhaps for these reasons, data streaming is only directly possible with Nextflow DSL-2 syntax, as of the time of writing (March 2021). For CWL, the language specification defines a ‘`streamable: true`’ field for output files, but direct support for this property is not yet part of the reference cwltool or other CWL runners. Neither Cromwell, Toil, nor Swift/T support piping either.

*Job retries* is an approach to retry a failed workflow step, and can be appropriate if the failure happened for an intermittent reason, such as service time-out, node unavailability or node failure. Swift/T allows retrying a failed job a number of times that can be set by the user, on the same MPI rank or on a randomly selected rank from those allocated to the workflow, possibly in other cluster nodes. Nextflow couples the `maxRetries`, `maxErrors`, and `errorStrategy` process directives to allow the user to `retry` a failed process, `ignore` the error, `finish` the run, or totally `terminate` the workflow effectively killing submitted processes. Similarly, Cromwell allows `maxRetries` as part of the `runtime` or `workflow options`, and also allows to either `ContinueWhilePossible` or resume but with `NoNewCalls` to quickly exit when a failure is detected.

*Caching or automatic check-pointing*, i.e., the ability to resume partial execution of a run, saves time and computational resources, especially when a large chunk of analysis has completed successfully. Swift/T is poor in this regard, as it always starts execution from the beginning, unless the programmer manually codes subworkflows separated at the anticipated checkpoints, and implements options to manually rerun subworkflows individually^17^. Nextflow is on the opposite end of the spectrum, permitting very granular access to workflow stages for restart purposes. This is accomplished by keeping all staged and intermediate files in a `work` directory with a `cache` directive enabled by default to index both scripts and input metadata (name, size, path, etc.). The granularity of restart can be controlled via `deep` and `lenient` modes. In contrast, call caching is by default disabled in Cromwell. It can be enabled by configuring a `MySQL` database instead of its default `HSQL` in-memory database and enabling the options for finer metadata checking, such as file hash caching, path prefix, and docker images’ tags.

### Debugging workflows

There are many reasons a workflow can fail: the nature of the data, a malfunction in the bioinformatics tool itself, improper setup or call of the bioinformatics tool, a bug in the execution engine, a hardware problem, an operating system issue, or something else entirely. Ideally, the log files for all these aspects would be cleanly separated, so that the workflow operator could easily trace the problem by hypothesizing the source and going through these logs one at a time. However, due to the asynchronous execution of independent workflow steps, messages can be echoed into the logs out of logical order, resulting in difficulty interpreting them. Each WfMS resolves this issue in its own way.

Swift/T provides a simple MPE-based model to track execution at the level of Swift and `turbine` operators. Messages are printed into a single log file in order of occurrence, not the order of the pipeline DAG, making it hard to discern invocations of individual bioinformatics tools. This makes it very difficult to determine the first step that failed and what data it was running on. Tool-level logs must be custom made, and even these logs can be difficult to interpret.

In contrast, as we mentioned before (section *Job execution: data staging*), Nextflow and the runners of CWL and WDL produce a canonical hierarchy of execution folders, with logs capturing the status of each workflow step saved into the same subfolder as the actual bash script being executed, along with the corresponding input and output data. Therefore, all the information about that particular step is in one place. In addition, the standard output from the executor is normally enough to establish which subfolder to inspect for signs of trouble.

### Monitoring the progress of workflow execution

The ability to monitor the progress of a workflow becomes critical with more tasks and increased workflow complexity. This monitoring facilitates scheduling, helps prepare the output data staging area, allows early detection of lag in a step, and yields information necessary for reporting, subsequent or retrospective analysis, and billing.

Nextflow supports several levels of detailed monitoring upon executing a workflow: (1) a crude `trace` report, (2) an html `timeline`, and (3) a complete `execution report`, including information about resources usage and processes runtime metrics (e.g., status, hash, command). Additionally, Nextflow is adding the Research Object (RO) model^43^, and thus adding greater transparency by uniquely identifying, collecting, and linking all provenance metadata of workflow runs^44^. Additionally, Nextflow has support for email notifications of workflow events like `onComplete` and `onError`, independent of the usual notifications from the CRM.

For WDL workflows, Cromwell only supports a `timeline` visualization, but only if run in `server` mode. The CWL community has developed `CWLProv`^45^, an informal profile standard defining how to record provenance of a workflow run as an RO using Linked Data standards^46^. This is implemented in the reference `cwltool`, and is planned for implementation in `toil-cwl-runner` too.

There are efforts to improve monitoring capabilities of these WfMS. Nextflow has the `Tower` platform (https://tower.nf/) for efficient monitoring and deployment; whereas WDL workflows can be submitted to a Cromwell server and examined via `cromshell`^47^ and Oliver^48^.

### Reproducibility and standardization

Reproducibility in biomedical analyses has become important recently^49,50^. For workflows, this means that anyone should be able to reconstruct the exact workflow run, including the correct sequence of steps, the actual commands, the runtime parameters and options, and the handling of data, e.g., chunking for parallelization, to reach the exact same conclusions despite differences in hardware, operating systems, and software dependencies. All this information must therefore be recorded in a way that is shareable and easy to understand, usually via code design documentation and the logs and RO described above.

Package managers, e.g., `Conda` and `Bioconda`^51^, provide means for clean shipping and installation of tools. Containerization technologies, e.g., `docker` and `singularity`, and their repositories (e.g `Dockstore`^52^ and `quay.io`) facilitate the reproducibility of computational pipelines. Both these advancements are increasingly integrated in recent releases of WfMSs. Nextflow utilizes a `conda` directive to specify packages needed by a given `process` and supports a `container` directive that allows `process`es to specify the `docker` or `singularity` images in which execution occurs. While lacking `conda` support, Cromwell can run tasks within a `docker` image specified in a WDL `task` or CWL `CommandLineTool` runtime options. `Singularity` images require special handling in the backend configuration file, but are supported too. Toil has similar features, but neither is supported in Swift/T.

Furthermore, the high complexity of biological workflows has driven the community to develop extra requirements for code documentation^21,42,52^. Users expect code annotation via extensive metadata including detailed workflow description, author information, and labels for stages, inputs, and outputs. Such metadata facilitate debugging, enhance overall project documentation via annotations on the DAG, help with maintaining and using a workflow written by someone else, and make code more searchable and citable. CWL and WDL each provide the code annotation capability via metadata blocks, and Nextflow via both the `manifest` scope of the configuration file and the directives of processes. CWL also includes support of EDAM ontology and SciCrunch identifiers for dependencies. Swift/T lacks this special workflow annotation feature, and only supports in-line comments.

Pragmatically, CWL recommended practices^21^ include shipping pipelines with permissive licenses and an SPDX identifier and using `SoftwareRequirement` to indicate dependencies and tool versions but warn against reliance on `InlineJavaRequirement` where possible. Adhering to these practices is a precursor to pipeline visualization and sharing via CWLViewer^38^. Likewise, curating Nextflow pipelines in nf-core^42^ requires using an MIT license, `docker` bundled software, stable release tags, a common pipeline structure, and continuous integration testing, in addition to passing their `nf-core lint` tests. They also recommend bundling the software via `bioconda`, using recent reference genome drafts and optimized output formats, and including a DOI, along with support and benchmarks from running in cloud environments^42^.

Taken together, the above efforts to enforce reproducibility of analyses have also resulted in a certain standardization of workflow implementation and distribution, which supports wider adoption of the WfMSs.

### Adoption and support

Nextflow, CWL and WDL are fruits of practicing bioinformaticians and computer scientists collaborating with biologists, who needed practical solutions to the problem of reliably performing their own large-scale analyses. Consequently, they enjoy greater adoption than Swift/T (table 3), albeit relying on comparably very permissive open source licenses. As a result, their evolution is rapid and highly community-driven, and user support is easy to find via mailing lists, gitter channels, Twitter, GitHub issues, etc. The community aspect is particularly important here, with numerous conferences, codefests, hackathons, and even GA4GH itself serving to develop, refine and cross-pollinate among the WfMSs^53–56^.

Adoption and support are further facilitated by commercial providers of genomics software-as-a-service. DNANexus provide `dxWDL` (https://github.com/dnanexus/dxWDL) and `dx-cwl` (https://github.com/dnanexus/dx-cwl). Seven Bridges developed `rabix` and many supporting utilities for CWL workflows, as they adopt CWL on their Cancer Genomics Cloud^57^. AWS provides the iGenomes database (https://registry.opendata.aws/aws-igenomes/) with the ability to directly use Nextflow and Cromwell on AWS Batch.

With this much collaborative cross-talk activity in the community, it is unsurprising that sometimes the boundaries between the WfMSs gets blurred. The ability to translate a workflow language into intermediate representation, e.g., the WOM representation of CWL and WDL code in Cromwell, enables *stitching meta-workflows* written in more than one language. It is quite possible that in the future some kind of a hybrid workflow coding paradigm might emerge, adopting the best from each of the current WfMSs and discarding the differences. Yet, a tricky aspect to this community-driven movement is that the evolution of a workflow language and its engine might sometimes outpace the development of a given pipeline. Therefore, conformance tests and backwards compatibility between a workflow language and an execution engine are critical.

### Cross-compatibility and conformance to standards

For a workflow language and an executor engine to work well together, they would ideally conform to the language specification and provide backwards compatibility to one another. This has not yet been widely the case, which is exactly what made this need apparent. For example, among WDL executors, both Toil and Cromwell can generate an abstract syntax tree from the same WDL code, except Toil’s `Hermes parser generator`^58^ translates tasks to Python functions, while Cromwell translates them into Bash. The supported code logic by each executor is also different: Cromwell does not allow nesting loops in version `draft-2` code, while Toil does not allow nesting a conditional within a loop or cascading tasks within a scatter body. In practice this leads to a lot of refactoring when switching from Cromwell to Toil. Similarly, different CWL runners support differing subsets of the possible `requirements` in the language specification or may even have different interpretations due to ambiguity in the language specification itself^36^. We believe the field would benefit from a wider conversation on this topic.

##### Highlights: Which WfMS to use day-to-day

In light of this, a pragmatic approach to workflow choice could be the following:

1. Assess: is there a need to build a new pipeline, or there is an existing reasonable pipeline in the Nextflow, CWL, or WDL repos?
  a. If a workflow exists that follows good coding practices, it should be adopted and modified as per specific needs.
  b. If starting fresh, without restrictions by collaborators’ preferences or existing legacy code-base:
    i. If a quick development cycle is important, Nextflow is optimal.
    ii. If code readability is important, WDL is optimal.
    iii. If execution environment is variable, or there is a need to work across heterogeneous hardware environments, CWL is optimal.
    iv. Table 1 is a quick overview of each language’s features at a crude level.
2. Assess: what execution constraints are in place?
  a. For HPC environments, pay particular attention to runners supporting differnt CRMs. Our recommended *free, production-scale* runners for these are: `Cromwell` (for both WDL and CWL), and `Nextflow` (for Nextflow workflows). Toil was less performant in comparison. (refer to section: *Scalability)*
  b. For running in the cloud, pay particular attention to runners with support for different cloud APIs, and features like automatic rescaling, containerization, and security settings.
  c. Table 2 is a quick overview of runners, language versions supported by each, and key performance aspects.

## Discussion

The choice of a WfMS for a specific use case is dependent on the immediate needs and resources of the application. Within the bioinformatics community, Nextflow, CWL, and WDL seem to be among the most adopted. The communities using and developing these three systems have been interacting closely since their introduction to the field, resulting in a very comparable set of semantic and engine features, though the nomenclature differs at times. Two other popular systems not examined in this study are Snakemake^59^ and Galaxy^60^. GA4GH TES support was only added to Snakemake in their November 2020 release, and `Galaxy` follows a rather different philosophy focusing on graphical user interface (while having its own CLI) and hence were both excluded.

To the same lines, this work has excluded other mature WfMS not GA4GH-supported like pegasus^61^, even though, at the time of writing, the pegasus team is developing utilities to import CWL workflows. Aside from arvados, this makes, pegasus an attractive first choice for running production-scale CWL code, compounding the project’s 20 years of experience in optimizing the performance features examined in this study, and uniquely having other characteristics like: 1) multitudes of interfaces, graphical and CLI-based, for real-time monitoring, debugging and reporting performance metrics; 2) ability to run seamlessly in heterogeneous grid, cloud and HPC staging sites; and 3) smooth data transfers for staging via many protocols, including `http, scp, GridFTP, iRods, AWS S3, … etc`^62,63^.

Another category is engines built as libraries in general-purpose programming languages, like Parsl^64^ (Python) and SciPipe^65^(Go). These engines give convenient access to the full expressiveness power and flexibility of the underlying language. Arguably, they are easier to learn, and hence, more attractive to adopt by a broader community than DSL workflow languages. At present, engines in this category seem less popular in the community though.

### Scientific and Business WfMSs

The emphasis on *Scientific* in this manuscript is to distinguish those WfMSs typically used in modeling and other scientific experiments, from those employed in business applications or other organizational contexts where human participants make decisions^7^. Accordingly, scientific workflows orchestrate tools (or services) based on data dependencies and often involve many data types. Business workflows on the other hand have a richer set of control flow constructs to support hierarchical dependencies^66^, logic correctness verification^67^, data flow modeling and consistency validation^68^, and resource requirement modeling and analysis^69^.

Additionally, the need to reuse and port workflows within the sciences is contrasted with restrictions on data and process access in business. Yet, the business community is more strict in developing and following standards^66^, with frequent evaluations and frameworks for benchmarking conformance with those standards^70,71^. The drive for standardization - despite early interoperability efforts, eg IWR^66^ and SHIWA^72^, has not surged in the sciences until recently, with ever larger scale collaborative and consortia projects^73,74^, and the push towards computational reproducibility^75^. Among others, this resulted in a heritage of bespoke systems, inconsistent terminology and inoperable formats, as a consequence of different WfMSs design requirements. Standardization, conformance evaluation and interoperability will continue to be an active area of research in scientific WfMSs.

### WfMSs in clinical and molecular diagnostic settings

Similar to research laboratories, clinical and diagnostic labs are concerned with the WfMSs aspects we examined above (also see supplementary note 1). However, there is focus towards properly developed, validated and operated pipelines that ensure the security of patient-identifying information, and the integrity and regulated access to data throughout each pipeline stage as per applicable laws and regulations^76,77^.

End-to-end validation using human samples, supplemented by in-silico validation, is both necessary and challenging given the constantly evolving and/or proprietary nature of computational tools, assay types and technology platforms. In fact, Roy *et al*. (2018) validation guidelines treat the bioinformatics pipeline as an integral part of the test procedure, and therefore require all its components to be validated, along with any filtering method applied to input data, and within an environment similar to the real-world lab where the pipeline will be used. Robust validation methods can involve the use of a *“golden”* set of workflow output files like `bam`s and `vcf`s based on human samples with known laboratory-validated variants. This way, concordance with known variants can be tested, and additions or modifications to the workflow made safely. This validation should be overseen by a qualified medical professional with NGS training, only after the complete pipeline has been designed, developed and optimized.

To allow for ongoing development without interfering with production-tested pipelines, it is beneficial to have separate stages in a clinical computing environment such as development, testing, and production. Code is developed and bugs are resolved in the development and testing stages, so by the time code gets to production, it has been robustly tested in the prior stages. Existing usable code in the production stage will not be changed or updated until all tests are passed. This allows the clinical labs to use the production code, and developers to push new code out simultaneously that will eventually be tested and deployed in production.

### Infrastructure as a predicate of WfMS design

A WfMS design is based on the infrastructure where it is intended to run. By employing an MPI library for parallel data communication, Swift/K^78^ for example, was made for large-scale computations on HPC environments extending to extreme scale supercomputing applications, and so are its successors - Swift/T^79,80^ and Parsal^64^- though Parsal has a wider bank of configurable executors including different cloud providers. On the other hand, other WfMSs like Toil^25^ and CWLairflow^81^ were developed for data analysis in the cloud, and hence supported containerization. Standard languages, CWL and WDL, aimed to enhance portability by obviating the need for intermediate data representation while allowing different groups to design and use executors that most fit their needs. This direct interplay between infrastructure and WfMSs will continue to play a key role in the design and composition of WfMSs well into the future with extreme scale systems and deep memory architectures^2^.

For example, while both *in situ* (i.e HPC) and *distributed* (i.e clouds and grids) workflows are challenged by analysis concurrency, locality and system topology awareness; more focus in the latter is paid towards security and crossing administrative domains^2^. Contrary, *in situ* designs, especially future exascale level, are challenged by power considerations, robustness, productivity with heterogeneous computing cores, increasingly complex hierarchical memory systems and small or no growth in bandwidth to external storage^2,82^. Consequently, Deelman *et al*.^2^ defines 4 key challenge areas for WfMS designs at the next scale: efficient task coupling, programming & usability, performance optimization & robustness, and validation & data integrity^83^- a list to which da Silva *et al*.^6^ add aspects like integrating big data analytics and human in the loop.

### Scalability

In bioinformatics, the community is still rather slow to adopt big data technologies, despite a few successful use cases (e.g., ADAM^84^, Gesal^85^, and most notably, the GATK’s move towards Spark re-implementations of existing trusted tools^26^). This is in part due to a need for rigorous and lengthy approval cycles for clinical applications^76^, and also more incentives and rewards for designing and building new tools rather than improving existing ones. Collectively, this means that in a majority of tools commonly used today, scalability has been thought of as an ad hoc- not as an integral part of software design. This manifests as more reliance on threading than MPI implementations for example, and an often complicated dependency stack for tools to work. Therefore, there is a real need for WfMSs that support complex execution patterns (at least DAGs) and large data volumes.

Yet, fairly benchmarking and reporting the scalability of different WfMSs remains elusive. For example, Swift/T papers demonstrate scalability in task throughput versus cores to extreme- and peta- scale computations^86,87^, congruent with the intent of its developers to use it for large-scale parallel applications. Similarly, Parsal literature differentiates *strong scalability*, running the same number of jobs (50,000) vs increased number of workers (up to 10^5^), from *weak scalability*, running the same number of tasks per worker (10) while increasing the number of workers^64^. Conversely, more bioinformatics-oriented WfMSs tend to demonstrate scalability in relation to the number or size of samples analyzed: Toil reports > 20,000 RNA seq samples analyzed on 1,000 nodes AWS `c3.8xlarge` cluster (each node of 32 cores, 60GB RAM, 640 GB SSD) in 4 days, and demonstrates scalability in terms of time and costs savings^25^, GLnexus quotes a 243,953 exome sequencing samples (33TB of compressed `gvcf`s) jointly called in 36 hours wall time with 1,600 threads^88^.

### Other comparative evaluations

Previous comparative manuscripts tend to be largely descriptive^2,6–9,89^. The closest to our work is Larsonneur *et al*.^16^, where testing was limited to a single node in a cluster, and examined WfMSs needed 4-6 minutes to run a genetic linkage analysis on variants from whole genome sequencing data of a trio^90^. This is a realistic bioinformatics use-case in terms of analysis complexity, but not in terms of computational requirements. It is also not clear how comparable the implementation was in the systems compared: Snakemake (python-based), Pegasus, Nextflow (java-based), Toil+CWL (python-based), and Cromwell+WDL (java-based). Regardless, they conclude reported performance differences are due to the algorithms used to compute job dependencies, and criticize java-based engines for consuming the most memory. Their results indicate that Pegasus, despite its very limited use in bioinoformatics, is optimal for HPC environments (performing better or at least similarly to the other top performing executors in most categories: elapsed time, CPU usage, memory, and number of inodes). Snakemake was best in terms of I/O wait time and idle time, and Cromwell did the best in terms of number of voluntary and involuntary context switches, but was the worst in most the other categories (closely followed by Toil). Therefore, they advocate for MPI-based execution engines.

A recent relevant paper is that of Jackson & Wallace^91^. By rapid prototyping, they quickly evaluated Snakemake, CWL+CWLtool, CWL+Toil, and Nextflow on a subset of RiboVis^92^ workflows. Like our approach, this gave them a better perspective than solely reading the documentation, tutorials or other review papers. Also, their criteria for selecting those WfMSs were adoption and support within the bioinformatics community, maturity, and licensing. However, the scope of their paper did not go into as great a depth in examining features as was done here.

### Trends & future directions

The revolution in the size and complexity of genomic data generation will continue to impact and be impacted by the progress in technology at the software and hardware levels. This manifests as a global trend in the community and funding bodies to attend to methods and software design, and also to plan for data analysis and handling as much as (if not more than) data generation. Another aspect are global efforts like those of the GA4GH and their designation of driver projects to refer to technology advancement at the levels of data transfers, security, storage and other relevant processing and accessibility aspects including ethics.

An optimistic trend for bioinformatics workflows is more attention being paid towards bridging the divide between user interface friendliness and expressiveness. On one hand, WfMSs like Galaxy that targeted user-friendliness from the beginning, are continuously expanding their code base with features to allow finer and more flexible control of execution details^60^. On the other, for the command-line frameworks examined here, many supporting tools exist that allow more friendly interfaces to the creation and deployment of workflow: NextflowWorkbench^27^ & DolphinNext^28^ for Nextflow; Rabix composer & `CWL-Experimental` (https://github.com/common-workflow-language/cwl-ex) for CWL; and `womtool` for WDL. Remarkably, away from `womtool`, all those supporting tools were contributed by the language’s broad community (and not its core developers’ team).

The growing need for portability of analyses also led to standard languages development^9^ - and ultimately decoupling the language specification from its engine implementation. Immediate benefits include better synergy between what a language offers and what patterns an analyst actually needs supported, and also more human readable (WDL) and/or machine interoperable (CWL) languages; while having open communications between communities supporting the different standards. Equally, executors supporting multiple backends, leading to GA4GH standards (like TES and WES- minimal APIs describing how a user submits a tool/workflow to an execution engine in a standardized way), and thereby giving more portability across platforms. For example, many runners have been developed to execute CWL code, including Rabix^93^, Avados, cwl-tes, Toil^25^, and AWE^94,95^; while WDL code can be run via Cromwell^26^ or Toil^25^. Additionally, there are runners for both languages on dedicated cloud platforms like DNAnexus, Seven Bridges, and Consonance. Concurrently, this encouraged other concerted efforts in areas like workflow provenance^96^, and design of more powerful graphical interfaces, like: Rabix Composer^93^ and CWLviewer^38^.

Thus, abiding by the FAIR principles^10^ for tools and workflows became a necessity as we move towards cloud computing^9^. For these purposes, many repositories exist today for sharing tools. Dockstore^52^, the standard implementation of the GA4GH TRS, is now hosting docker-based tools described in Nextflow, besides CWL and WDL. Projects like bio.tools (https://bio.tools/), an ELIXIR Tools & Data Services Registry, support the findability and interoperability of bioinformatics application software by employing biotoolsSchema and EDAM ontologies^97^ for software description and annotation. BioContainers^98^ is also another remarkable project as it hosts both both docker and rkt images, with special focus on tools in proteomics, genomics, transcriptomics and metabolomics. The ability of a WfMS to seamlessly fetch images from these repositories or publish workflows will undeniably be a bonus in today’s market; along with the core features of running in heterogeneous environments, robustness and scalability. As the field evolves, the need for systematic performance benchmarking^70^ and conformance testing frameworks^71^ grows.

## Methods

### Nomenclature

We use the terms *workflow* and *pipeline* interchangeably to describe the steps of a given analysis or its implementation. An analysis *step*, or *stage*, of a workflow, means a self-contained computational unit, including serial invocation of multiple command line tools. In some WfMS nomenclatures this is called a *process* (in `Nextflow`), *task* (in `WDL`), or *leaf app* (in Swift/T). A workflow is therefore a collection of these steps strung together according to some logic, though a workflow script may consist of *sub-workflows*, a group of steps that are themselves a workflow. It is the *execution engine*, *runner*, or *executor* that handles the execution details of running these stages. Finally, we use the word *job* for whatever computation is submitted to the resource manager in an HPC setting.

### Use case I: variant calling pipeline

Genome Analysis Toolkit (GATK)’s variant calling pipeline^99,100^ is a typical bioinformatics pipeline that is ideal for testing the expressiveness of a workflow language and the capabilities of a workflow execution engine (Fig Supplementary 1). To curtail lengthy run times, proper implementation of this pipeline requires packaging calls to the bioinformatics tools and interleaving the phases of serial and parallel processing over multiple samples. Care must be taken to avoid overwhelming the HPC cluster with a large number of jobs or overusing the compute resources. Additionally, upholding the maintenance, portability, and reproducibility of this pipeline can be taxing on the developer, due to its many steps and large number of configurable parameters (supplementary note 1). We used a synthetic dataset, based on the `hg19` human genome assembly and using Illumina TrueSeq v1.2 targeted regions (Whole Exome Sequencing) to 30X sequencing depth, generated by the NEAT simulator^101^. Simulation code is linked to in section *Additional information*.

### Use case II: scalability evaluation

Scalability is an important feature of a WfMS that reflects its ability to parallelize across an increasing number of tasks without slowing down processing. To test the scalability of WfMSs without biasing the results with the intricacies of variant calling, we built a simplified workflow: Each task was simply a `hostname` command, scattered as an array of `n` parallel processes across `n` cores (i.e., a *1-step workflow*). Additionally, we tested how task dependency chains affected scalability by building a cascade of two identical tasks, also scattered `n` times across `n` cores (i.e., a *2-step workflow*, Fig 4). We performed multiple runs while varying the scalability parameter `n` from 1 to the maximum number of cores in the cluster, and tracked which node every task was deployed on and when. This allowed examination of how well the tasks were distributed among the nodes, the overhead to initiate the WfMS, and the maximum possible throughput.

**Figure 4.**
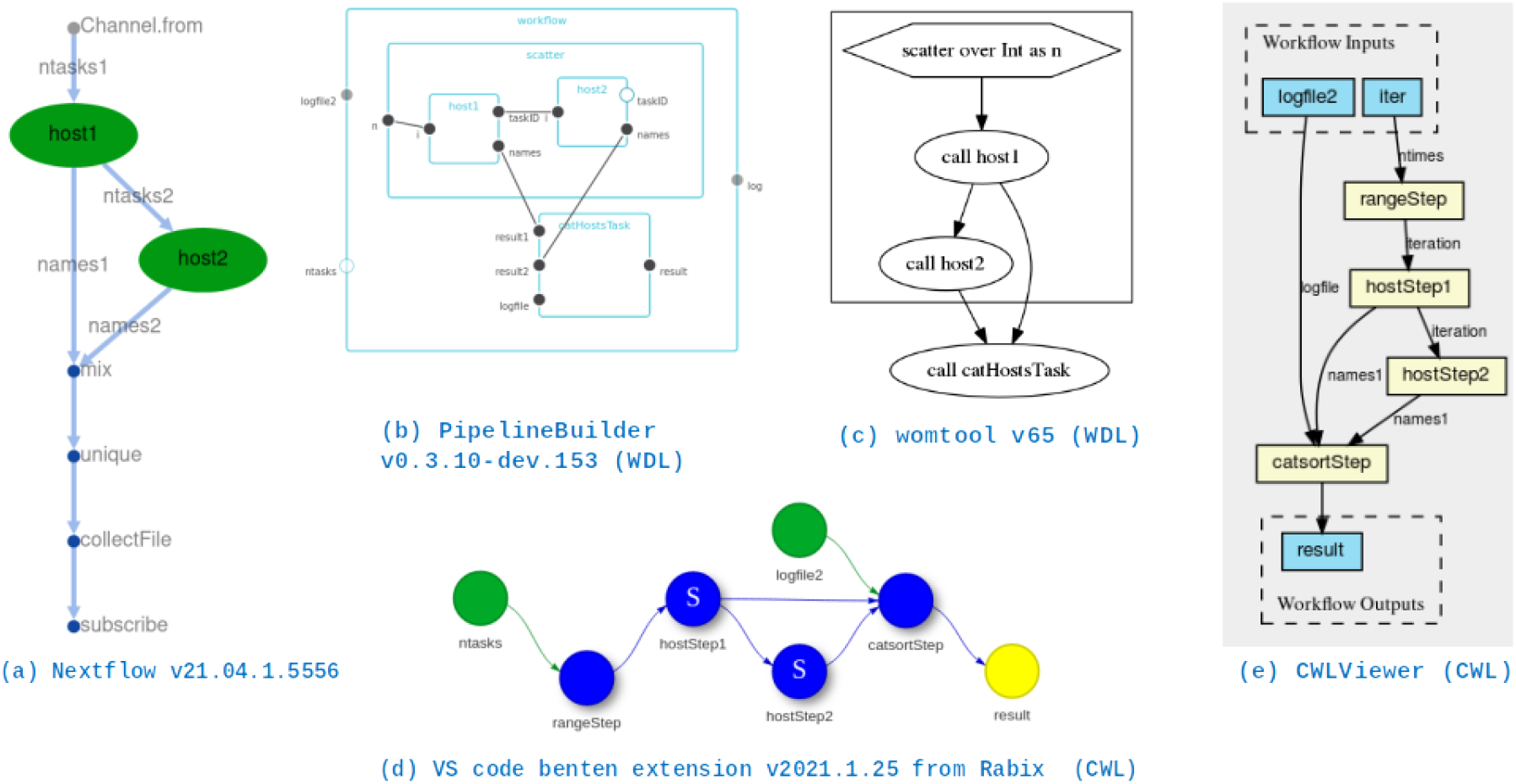
DAGs corresponding to a simple workflow of 2 processes (besides output aggregation) used to assess the scalability of the executors of section *Scalability*, as generated by the most recent version of each executor or utility visualizer of each language in July 2021

### WfMSs under consideration

We chose WfMSs (Table 4) that support the widely adopted Global Alliance for Genomics and Health (GA4GH) APIs: the `Workflow Execution Service Schema (WES)` for describing how a user submits a workflow to an execution engine in a standardized way, the `Task Execution Schema (TES)` for describing batch execution tasks (implemented primarily as `Funnel:` https://github.com/ohsu-comp-bio/funnel), and the `Tool Registry Service (TRS)` for sharing code (implemented primarily as `dockstore:` https://dockstore.org). These WfMS languages are Common Workflow Languagen (CWL)^102^, Workflow Description Language (WDL)^26^ and Nextflow^103^. For contrast, we included Swift/T^79,87^, a WfMS developed and used primarily in peta- and exascale physics applications^86^. We evaluated the language properties of Swift/T, but did not test its scalability as it has been thoroughly examined earlier (e.g.,^82,87^). All four of these WfMSs use the *dataflow* paradigm to provide implicit parallelism in running computations based on the availability of data and compute resources^31^, making them suitable for our use case of massively high-throughput production environments.

### Computational systems

The above pipelines were developed on personal computers, then ported to an HPC machine (Biocluster)^104^ and to Amazon cloud AWS (Table 5). Biocluster utilizes Slurm for job scheduling and includes five `Supermicro SYS-2049U-TR4` nodes of 72 Intel Xeon Gold 6150 2.7 GHz cores, 1.2 TB RAM each. In AWS we tested three different environments: (1) `Batch`, provided by AWS, for optimal provisioning of compute resources during batch processing; (2) `Cloud cluster`, which is built by Nextflow at run time and uses `Apache Ignite` for resource management; and (3) a dedicated, fixed-size Slurm `Parallel cluster` that we constructed out of 100 worker nodes and a head node, all `m5a.24xlarge` instances with 96 cores and ~412 GB RAM each, configured using default AWS settings.

## Acronyms

API: Application Programming Interface
AWS: Amazon Web Services
CLI: Command Line Interface
CPU: Central Processing Unit
CRM: cluster resource manager
CWL: Common Workflow Languagen
DAG: Directed Acyclic Graph
DOI: Digital Object Identifier
DSL: Domain Specific Language
EDAM: EMBRACE Data And Methods, a bioinformatics (dry) ontology
FAIR: Findable, Accessible, Interoperable and Reproducible
GA4GH: Global Alliance for Genomics and Health
GATK: Genome Analysis Toolkit
GCP: Google Cloud Platform
GUI: Graphical User Interface
HPC: High Performance Computing
IWR: Intermediate Workflow Representation
JVM: Java Virtual Machine
MPE: MPI Parallel Environment
MPI: Message Passing Interface
NGS: Next Generation Sequencing
RAM: Random access Memory
RO: Research Object
SHIWA: Sharing Interoperable Workflows for large-scale scientific simulations on Available DCIs
Tcl: Tool Command Language
TES: Task Execution Schema
TRS: Tool Registry Service
WDL: Workflow Description Language
WES: Workflow Execution Service Schema
WfMS: Workflow Management System
WOM: Workflow Object Model

## Acknowledgements

This work was a product of the Mayo Clinic and Illinois Alliance for Technology-Based Healthcare. Special thanks for the funding provided by the Mayo Clinic Center for Individualized Medicine and the Todd and Karen Wanek Program for Hypoplastic Left Heart Syndrome. We also thank the Interdisciplinary Health Sciences Institute, the Carl R. Woese Institute for Genomic Biology and the National Center for Supercomputing Applications for their generous support and access to resources. We particularly acknowledge the support of Keith Stewart, M.B., Ch.B., Mayo Clinic/Illinois Grand Challenge Sponsor and Director of the Mayo Clinic Center for Individualized Medicine. Special gratitude to Gay Reed and Amy Weckle for managing the project. Many thanks to the Biocluster team for their consultation and advice during the deployment of our workflows on their machine. Thanks also to the UIUC AWS infrastructure, and the AWS Research Credits Award for supporting this work. Finally we are grateful for the support of H3ABioNet, funded by the National Institutes of Health Common Fund under grant number U41HG006941.

## Author contributions statement

- *Conceptualization*: AEA, SNH, MEH, EWK, NM, FMF, LSM
- *Data curation*: AEA, TB, MEH, LSM
- *Formal analysis*: AEA, LSM
- *Funding acquisition*: MEH, LSM
- *Investigation*: AEA, TB, FMF, LSM
- *Methodology*: AEA, NM, RV, FMF, LSM
- *Project administration*: SNH, MEH, GDK, EWK, KIK, FMF, LSM
- *Resources*: FMF, LSM
- *Software*: AEA, JMA,TB, PB, JRH, DDI, MCK, MTK, CAR, LSM
- *Supervision*: MEH, GDK, EWK, SMS, FMF, LSM
- *Validation*: AEA, TB, NM, LSM
- *Visualization*: AEA, JMA, LSM
- *Writing-original draft*: AEA, JMA, CEF, MCK, FMF, LSM
- *Writing-review and editing*: AEA, JMA, CEF, SNH, MEH, MCK, FMF, LSM

## Additional information

### Data and code availability

The synthetic WES dataset used for the performance analysis of variant calling workflow will be made available by the authors, without undue reservation, to any qualified researcher. The commands used to generate this synthetic data are available at https://github.com/ncsa/MayomicsVC/tree/dev-gatk. Other code and data are provided in the respective repositories of table 4.

