## Supplementary notes and figures for "Design considerations for workflow management systems use in production genomics research and the clinic"

#### Supplementary materials

### Contents

**1 Design considerations in building the variant calling pipeline** **2**

**2 Workflows Invocations** **3**

**3 Coding examples** **4**

**4 Working directory structure** **5**

†Current address: Center for Computational Biology, University of California, Berkeley, CA, USA
‡These authors equally supervised this work

### 1 Design considerations in building the variant calling pipeline

Design considerations for building a Swift/T-defined Variant Calling pipeline are detailed elsewhere [2], so in this section, we only focus on respecting *Modularity* with an architecture that allows consistent evaluation of Swift/T along with the other 3 WfMSs: Nextflow, CWL and WDL.

In this context, modularity means the ability to construct a complete `workflow` from a set of smaller and independent `processes`, `apps`, `CommandLineTools`, or `tasks`/`subworkflows` (as per the semantics of each WfMS (see section *Methods: Nomenclature*)).

To meet the modularity constraint, `src` code is arranged as per Fig Supplementary 2a into folders corresponding to each WfMS language, and within which there are folders for calling tasks, unit-testing those tasks, and defining the logic of workflows composed of these tasks (except for Nextflow). The tasks themselves were written in conditionals-free, stand alone bash scripts that provide consistent output definitions and logging functionality regardless of inputs specifications and bioinformatics tools being called (Fig Supplementary 2b). These bash scripts are also free from streaming (i.e., piping) between processes for more robustness and easier debugging of failure source when needed. This further allows seamless switching between Sentieon-based [3] and GATK-based [4, 5] tools (or others) while using the same WfMS (and vice-versa). For easier working with input `json` files (in case of WDL- and CWL- defined pipelines), helper parser and validator scripts were written in python to populate values from an easy to construct configuration file into the needed input `json`. This added a layer of abstraction/independence between the processing logic of the workflow (conditions and loops- the DAG definition), and the underlying invocations of the bioinformatics tools. Additionally, it allowed a head-to-head comparison between the 3 languages (See sections *Results: Language expressiveness* - *Results: Support for modularity*).

The 4 chosen WfMSs considered here (Nextflow, Swift/T, CWL and WDL) all have engines that adopt the *dataflow*-paradigm. This means an inherent and implicit parallelism in running computations based on data (and resources) availability (rather than location within a script)- making them appropriate for sprouting parallel jobs rather easily (compared with, say, native

**Pseudo code of a multi-sample variant calling pipeline**

```
for each sample:
   for each LANE:
      if (pairedEndReads?): PairedEndAlignment
      if (singleEndReads?): SingleEndAlignment
   Merge aligned bams
   if (deduplicate?): Deduplication
   for each CHR:
      Base Quality Score Recalibration
      Haplotyper
   Merge gvcfs
for each CHR:
   Joint Calling
GatherVCFs
```

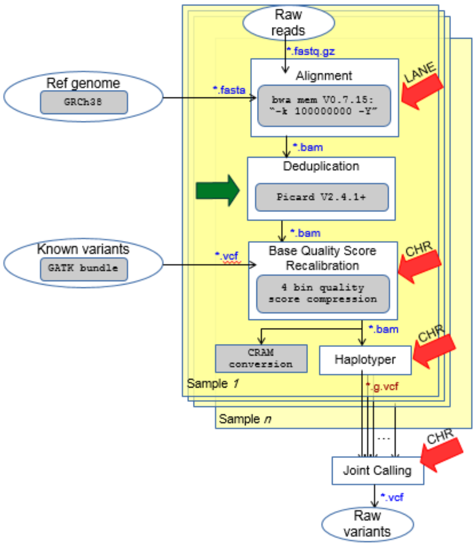

**Figure Supplementary 1: Analysis stages in a typical Genome Analysis Toolkit (GATK)-based multi-sample Variant Calling pipeline where each yellow slice is a sample. Gray blocks denote functional equivalence recommendations [1]- with re-alignment (after Deduplication) not shown. Red arrows denote parallel stages, and Green arrows denote optional stages, and thus need a WfMS that supports:Analysis sequential, parallel (looping) and conditional processing and also nesting these within an overall loop.**

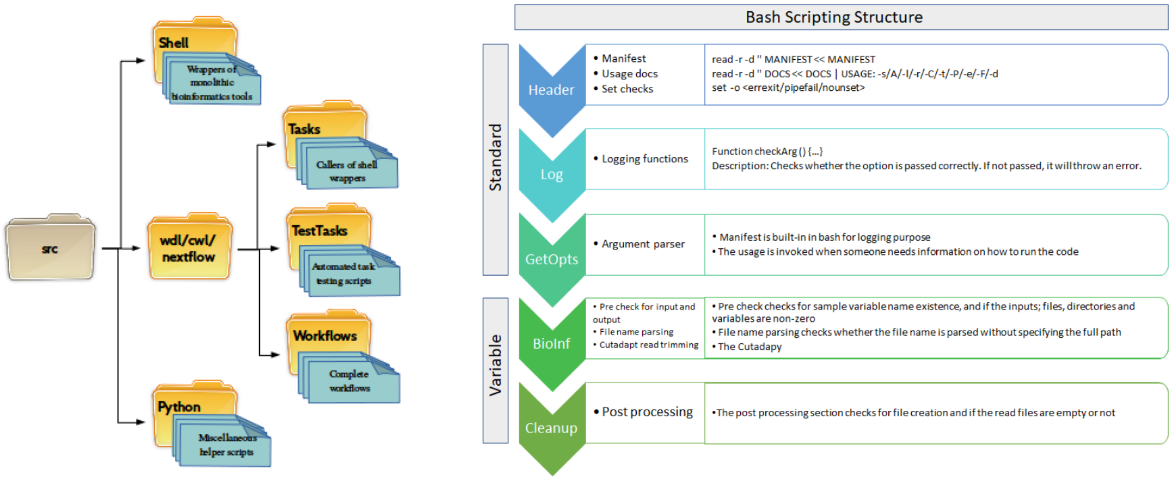

**(a) Code directory structure**

**(b) Shell script design**

**Figure Supplementary 2: Architectural code organization of the implemented variant calling pipeline in WDL and Nextflow. A slightly different organization was followed for the Swift/T repo, but it follows similar ideas**

bash parallelization and other top-down sequential languages). A complete evaluation of these parallelization and run-time features follows in the main results sections (*Data dependencies and parallelism* - *Workflow dependency graph resolution and visualization*). Performance aspects are discussed in the remaining results sections (*Executor-level differences* - *Robustness*). Operational aspects, like debugging and cross-compatibility are examined in the remaining results sections (*Debugging workflows* - *Cross-compatibility and conformance to standards*).

### 2 Workflows Invocations

**Custom Nextflow invocation**

```
$ nextflow run workflow.nf -c backend_runtime_and_input.conf
```

**Custom Swift/T invocation**

```
$ swift/t -m backend_name -s runtime_conf.sh workflow.swift
```

**Custom Cromwell invocation**

```
$ # with WDL
$ java -Dconfig.file=backend_and_runtime.conf -jar cromwell.jar run workflow.wdl --
    inputs inputs.json --options workflow_options.json
$ # with CWL
$ java -Dconfig.file=backend_and_runtime.conf -jar cromwell.jar run workflow.cwl --
    inputs inputs.yml --type cwl --options workflow_options.json #The --options
    workflow_options.json directive was not respected in our tests in Jan 2020
```

**Figure Supplementary 3: Typical invocations of workflows written in each language examined in this study, with configuration options specifying the backend against which to run the workflow, its runtime settings and inputs to the workflow run. Example configuration files are provided in our `scalability-tst` repo.**

### 3 Coding examples

#### Nextflow DSL-1 process

```
// Defining and using a process (in Tasks/alignment.nf):
SampleNamesChannel = Channel.from(params.SampleName.tokenize(','))  // Comma-separated list of strings input
process Alignment {
  input:
  val SampleName from SampleNamesChannel // Implicit Parallelism over channel elements
  """
  /bin/bash alignment.sh ...            # Script in the 'shell' directory
  """
}
```

#### Swift/T leaf function and its usage

```
// Defining a leaf function (in bioapps/align_dedup.swift):
@dispatch=WORKER
app (<outputs>) bwa_mem (<inputs>) {<command invocation>}
app (<outputs>) samtools_view (<inputs>) {<command invocation>}
```

```
// Using the leaf funciton:
import bioapps.align_dedup;
foreach sampleName in samples {            // Explicit parallelism over samples
  string AlignDir = dircat(vars["OUTPUTDIR"], sampleName);
  mkdir(AlignDir) =>
  alignedsam = bwa_mem(sampleName);        // Explicit dependency via '=>' operator
  alignedbam = samtools_view(alignedsam); // Implicit dependency via inputs
}
```

#### CWL v1.0 CommandLineTool and workflow

```
# Defining a CommandLineTool (as 'alignmentCommand.cwl'). 'dedupCommand.cwl' is defined similarly:
cwlVersion: v1.0
class: CommandLineTool
baseCommand: alignment.sh  # Script in the  'shell' directory
inputs:
  sampleName:
    type: string
outputs:
  alignedbam:
    type: File
    outputBinding:
      glob: '*.bam'
```

```
cwlVersion: v1.0
class: Workflow
requirements:
  ScatterFeatureRequirement: {} # For parallelization over input array items
inputs:
  samples_array: String[ ]
outputs: [ ]
steps:
  alignmentStep:
    run: alignmentCommand.cwl
    scatter: sampleName     #  Explicit Parallelism over samples
    in:
      sampleName: samples_array
    out: [alignedbam]      # Step output
  dedupStep:
    run: dedupCommand.cwl  # CommandLineTool definition not provided, but similar to alignmentCommand.cwl above
    scatter: sampleName
    in:
      sampleName: alignmentStep/alignedbam # Implicit dependency via inputs
    out: [ ]
```

#### WDL v1.0 task and workflow

```
# Defining a task (in Tasks/alignment.wdl):
version 1.0 #if omitted, defaults to version "draft-2"
task alignmentTask {
  input {
    String SampleName
  }
  command {
    /bin/bash alignment.sh ...  // Script in the 'shell' directory
  }
}
```

```
version 1.0 #if omitted, defaults to version "draft-2"
import "Tasks/alignment.wdl" as ALIGN
workflow RunAlignmentTask {
  scatter (sampleName in samples) {              # Explicit Parallelism over samples
    call ALIGN.alignmentTask as ALIGN_paired     # Task defined with output `alignedbam`
    call DEDUP.dedupTask as dedup {input: ALIGN_paired.alignedam}   # Explicit dependency via inputs
  }
}
```

**Figure Supplementary 4: Minimal examples demonstrating equivalent parallel and sequential tasks within a variant calling pipeline in each WfMS language. The relative length and sophistication of CWL code can be appreciated here**

### 4 Working directory structure

The workflow used here is the 1-step version of the pipeline used for testing scalability. There is a `hostname` process that is run twice in parallel, then unique hostnames are sorted and collated in a file. Testing was done in a local machine, and relevant comments accompany each workflow run. The complete code can be found in our `scalability-tst` repo here: https://github.com/azzaea/scalability-tst

#### 4.1 Nextflow

`Nextflow` defaults to creating a `work` directory where it is run. Each process will have its own hexa-coded directory of all inputs, outputs, intermediates and logs. No need for a dedicated `cat` process with `Nextflow`, since it has efficient `channel` operators for organizing such outputs.

The output of the workflow is sent to a specific directory in our code. Its contents are shown in the snippet below.

**Nextflow invocation**

```
$nextflow -version

N E X T F L O W
version 19.10.0 build 5170
created 21-10-2019 15:07 UTC (17:07 CEST)
cite doi:10.1038/nbt.3820
http://nextflow.io

$
$nextflow run host_process.nf -profile standard --ntasks=2 --log=log.txt
## log omitted
$
$ cat results.nf/hosts/log.txt
azza-Satellite-P845
$
$ tree work
```

```
work/
├── 29
│   └── 2fd42e9c4edfc45980bd3dac003c9b
│       ├── .command.out
│       ├── .command.sh
│       ├── .command.begin
│       ├── .command.log
│       ├── .command.err
│       ├── .command.run
│       └── .exitcode
└── 1b
    └── 31200e421cdb1ab2bee26d5460147c
        ├── .command.out
        ├── .command.sh
        ├── .command.begin
        ├── .command.log
        ├── .command.err
        ├── .command.run
        └── .exitcode
```

4 directories, 0 files

#### 4.2 WDL: Cromwell

`Cromwell` defaults to creating a `cromwell-execution` directory where it is run. Each workflow will have its own directory, and different runs will be different hexa-coded subfolders within. Tasks will further have their own directories nested within their parent sub-workflows or scattering patteren- if present. Similar to Nextflow, each process directory will host all of its inputs, outputs, intermediates and logs.

The output of the workflow is sent to a specific directory via the `workflow.options.json` file. Its contents are shown in the snippet below.

**cromwell invocation**

```
$ java -jar $crom --version
cromwell 42
$
$ java -jar cromwell-42.jar run host_process.wdl --inputs host_process_workflow.json --
    options  workflow_options.json
## log omitted, containing final outputs location within cromwell-executions dir before
    they are copied to destination specified within workflow.options.json
$
$ cat results.cromwell/hosts/log.txt
azza-Satellite-P845
$
$ tree cromwell-executions
```

```
cromwell-executions/
└── f247f741-f15b-4ff7-b661-2b87a9121fd1
    ├── call-catHostsTask
    │   ├── tmp.246499fd
    │   └── execution
    │       ├── script.submit
    │       ├── script
    │       ├── log.txt
    │       ├── rc
    │       ├── stderr
    │       ├── stderr.background
    │       ├── stdout
    │       ├── stdout.background
    │       └── script.background
    └── call-host1
        ├── shard-1
        │   ├── tmp.0a8ca919
        │   └── execution
        │       ├── script.submit
        │       ├── script
        │       ├── rc
        │       ├── stderr
        │       ├── stderr.background
        │       ├── stdout
        │       ├── stdout.background
        │       └── script.background
        └── shard-0
            ├── tmp.30a11e11
            └── execution
                ├── script.submit
                ├── script
                └── rc
```

```
│
├── stderr
├── stderr.background
├── stdout
├── stdout.background
└── script.background
```
12 directories, 25 files

##### 4.3 WDL: toil-wdl-runner

`toil-wdl-runner`: defaults to deleting the working directory, and does not understand command line arguments acceptable otherwise to Toil. Hence, below we explicitly generate a python equivalent of our WDL code and edit it to accept command line options for specifying a working directory and not deleting it upon successful workflow run.

Additionally, `Toil` doesn't seem to have the ability to put output files in a user desired destination. Instead, it puts them in the current directory from which it is run. The hostnames retrieved in this case are unusual- preceded by apostrophe or letter (b).

**Toil invocation**

```
$ toil --version
4.1.0
$
$ toil-wdl-runner --dev_mode 3 host_process.wdl host_process_workflow.json
## This mode translates our wdl code into python and produces a file named:
    toilwdl_compiled.py
$
$ sed -i 's/.*getDefaultOptions.*/    parser = Job.Runner.getDefaultArgumentParser()/'
    toilwdl_compiled.py
## To allow passing command line options
$
$ sed -i 's/.*options.clean.*/    options = parser.parse_args()/' toilwdl_compiled.py
## To prevent deleting the working directory
$
$ mkdir workDir
$
$ python toilwdl_compiled.py --workDir workDir --cleanWorkDir never myJobStore
## log omitted, no pointers to where outputs are; but they are placed in this directory
$
$ cat log.txt
'
b'azza-Satellite-P845
$ tree workDir
```

```
workDir/
└── node-991a1f3e-c498-44af-98a0-ce3b4698291c-2a77c9e44cbe4a17b74b11479a5c5836
    ├── tmprfhl6a73
    │   ├── worker_log.txt
    │   └── 697be4c6-1294-4982-a484-9408ae2b00fc
    │       ├── log.txt
    │       └── tj9id25at
    │           └── execution
    ├── tmpv3sjqcun
    │   ├── worker_log.txt
    │   └── efe248ad-e14b-466b-9331-225fe59e2e07
    ├── tmpc8ux75re
    │   ├── worker_log.txt
    │   └── 62c036a4-5252-413f-b084-61ea8711a532
    │       └── t8hk6ry_n
    │           └── execution
    └── tmp_2o9cnwe
```

```
│   ├── worker_log.txt
│   ├── 569830e3-645c-4749-87d9-a5b017bfaaa4
│   │   └── tpoe1tqu9
│   │       └── execution
│   ├── 66b4264f-93fd-4469-acf9-2511b72da37b
│   ├── 39c81999-e94e-4383-918e-3941352eac67
│   ├── 16b55c7b-8a47-4725-afed-c35ad581f7e3
│   └── 9f67794f-127b-4f22-981f-791e2af310aa
├── tmppiv6fdmx
│   └── worker_log.txt
├── tmpab7pga6d
│   ├── worker_log.txt
│   └── f5634d44-86cc-4898-b10e-54b31c48806a
│       └── telqlwbjb
│           └── execution
└── tmploar_wsi
    └── worker_log.txt
```

25 directories, 8 files

##### 4.4 WDL: miniWDL

`miniWDL` defaults to creating a timestamped named working directory per each workflow run, appended by the workflow name. It requires that only inputs that the workflow actually uses are present in the input json file. Under the hood, for miniWDL to run locally, docker needs to be installed with proper user permissions. A parallelized workflow will consequently be run in `Docker swarm` mode. This explains the ouput in the example below- hostnames are from this docker swarm environment (not the local environment).

Similar to `Toil`, `miniWDL` does not offer the possibility to place outputs in a user defined destination. It doesn't place outputs in the current directory either, but the execution log will direct to their location within the execution directory.

**miniWDL invocation**

```
$ miniwdl --version
miniwdl v0.7.4
miniwdl.plugin.file_download    gs = WDL.runtime.download:gsutil_downloader     miniwdl
    0.7.4
Cromwell 47
$
$ miniwdl run -i host_process_workflow.json host_process.wdl
## log omitted, containing final outputs location within the timestamped execution
    directory
$
$ cat /home/azza/github_repos/varCall/scalability-tst/src/wdl/20200604_143731_hostwf/
    output_links/log/log.txt
8ef5b47e1ee3
af718034688c
$
$ tree 20200604_143731\_hostwf
```

```
20200604_143731_hostwf/
├── inputs.json
├── outputs.json
├── rerun
├── workflow.log
└── call-host1-1
    ├── task.log
    └── inputs.json
```

```
│
│   ├── outputs.json
│   ├── command
│   ├── stderr.txt
│   ├── stdout.txt
│   ├── work
│   └── output_links
├── call-catHostsTask
│   ├── task.log
│   ├── inputs.json
│   ├── outputs.json
│   ├── command
│   ├── stderr.txt
│   ├── stdout.txt
│   ├── work
│   │   └── log.txt
│   └── output_links
│       └── result
│           └── log.txt
├── wdl
│   └── host_process.wdl
├── output_links
│   └── log
│       └── log.txt
└── call-host1-0
    ├── task.log
    ├── inputs.json
    ├── outputs.json
    ├── command
    ├── stderr.txt
    ├── stdout.txt
    ├── work
    └── output_links
```

16 directories, 26 files

#### 4.5 CWL: cwltool

`cwltool` does not create a working directory, and outputs are placed directly in the current directory.

**cwltool invocation**

```
$ cwltool --version
/home/azza/pythonenvs/toil3/bin/cwltool 3.0.20200324120055
$
$ cwltool host_process.cwl host_process_workflow.yml
## log omitted, containing final outputs and their locations
$
$ cat /home/azza/github_repos/varCall/scalability-tst/src/cwl/log.txt
azza-Satellite-P845
```

#### 4.6 CWL: Cromwell

The general notes of section 4.2 apply here, except that the `-options` directive is not respeced by Cromwell, and hence there is no way to specify the final destination of output files readily.

Instead, the log gives complete path to where outputs are stored within the `cromwell-execution` directory

**Cromwell invocation**

```
$ java -jar $crom --version
cromwell 42
$
$ java -jar $crom run host_process.cwl -i host_process_workflow.yml --type cwl
## log omitted, containing final outputs location within cromwell-executions dir
$
$ cat /home/azza/github_repos/varCall/scalability-tst/src/cwl/cromwell-executions/
    host_process.cwl/b13c231f-b3aa-4880-b503-afc0edf541e8/call-catsortStep/execution/log
    .txt
azza-Satellite-P845
$
$ tree cromwell-executions
```

```
cromwell-executions/
└── host_process.cwl
    └── b13c231f-b3aa-4880-b503-afc0edf541e8
        ├── call-catsortStep
        │   ├── inputs
        │   │   ├── 1264064947
        │   │   │   └── result.host.txt
        │   │   └── -532886412
        │   │       └── result.host.txt
        │   ├── tmp.dd2a99cc
        │   └── execution
        │       ├── script.submit
        │       ├── glob-b34dfc006a981a93d6da067cf50036fe.list
        │       ├── script
        │       ├── log.txt
        │       ├── rc
        │       ├── stderr
        │       ├── stderr.background
        │       ├── log.txt.background
        │       ├── script.background
        │       └── glob-b34dfc006a981a93d6da067cf50036fe
        │           └── cromwell_glob_control_file
        └── call-hostStep1
            ├── shard-1
            │   ├── tmp.3bf79c99
            │   └── execution
            │       ├── script.submit
            │       ├── glob-b34dfc006a981a93d6da067cf50036fe.list
            │       ├── script
            │       ├── rc
            │       ├── stderr
            │       ├── stderr.background
            │       ├── result.host.txt
            │       ├── result.host.txt.background
            │       ├── script.background
            │       └── glob-b34dfc006a981a93d6da067cf50036fe
            │           └── cromwell_glob_control_file
            └── shard-0
                └── tmp.1637f792
```

```
        └── execution
            ├── script.submit
            ├── glob-b34dfc006a981a93d6da067cf50036fe.list
            ├── script
            ├── rc
            ├── stderr
            ├── stderr.background
            ├── result.host.txt
            ├── result.host.txt.background
            ├── script.background
            └── glob-b34dfc006a981a93d6da067cf50036fe
                └── cromwell_glob_control_file
└── call-rangeStep
    ├── tmp.01de7c14
    └── execution
        ├── script.submit
        ├── glob-b34dfc006a981a93d6da067cf50036fe.list
        ├── script
        ├── rc
        ├── stderr
        ├── stderr.background
        ├── stdout
        ├── stdout.background
        ├── script.background
        └── glob-b34dfc006a981a93d6da067cf50036fe
            └── cromwell_glob_control_file
```

22 directories, 42 files

#### 4.7 CWL: toil-cwl-runner

The general notes of section 4.3 apply here, except that `toil-cwl-runner` accepts command line options directly.

**Toil invocation**

```
$ toil --version
4.1.0
$
$ mkdir workDir
$
$ toil-cwl-runner --workDir workDir --cleanWorkDir never host_process.cwl
    host_process_workflow.yml
# log omitted
$
$ cat log.txt
azza-Satellite-P845
$
$ tree workDir
```

```
workDir/
└── node-89dc310f-9595-4a9e-97aa-f71d3f24652d-2a77c9e44cbe4a17b74b11479a5c5836
    └── tmp6s3gi8fm
        ├── worker_log.txt
        └── 1a683141-6c14-4ab1-bbc4-b093246bb5bf
            ├── tzxrehcmg
            │   └── out
            └── t8syjkal3
```

```
            tmp-outsm57wg9r
            tmp-outbmlgcskr
        tah5qtjw0
        tnq9ijiwb
    7c55fea3-7e03-495d-9636-ce8af16bbf80
tmpymlh10pb
    worker_log.txt
tmpi64silti
    worker_log.txt
    7222d661-95b0-445b-be5b-a12b83ca1aa8
tmph9k9tuhb
    worker_log.txt
tmpks6eh_0m
    worker_log.txt
tmpzek_gzg3
    worker_log.txt
    79905428-016a-4cd1-b968-95cb5bbfa001
        txbku8b2q
        t231lba6i
            out
        tl4zpjmr7
            tmp-out2774dp3e
            tmp-outn66yom55
                result.host.txt
        tqix7a77y
tmpgme2si7h
    worker_log.txt
    c9f654f1-5167-422e-b913-e6ff8aa575fb
tmpn38udpjq
    worker_log.txt
    d023bfcb-e40e-48bf-af3f-a26b347ae3e0
        ttcurwhla
        tvjhie4dj
            tmp-out7a0iq94w
                result.host.txt
            tmp-outu6c68t67
        t81_96enr
            out
        tchmo684i
tmpm0l60nc6
    worker_log.txt
    54da9631-5317-46f3-8965-4cbb6f842b70
tmp1s19t2nm
    worker_log.txt
    b2ab1ed4-247a-44ac-82c0-f6ea7584c9a0
        tmphzzn5mh2.tmp
        tmp4c_nsumn.tmp
        tl17es875
            out
        taqvgxqja
        tz_walc9l
```

```
└── t_g6ohtq5
    ├── tmp-out1coaxjj6
    │   └── log.txt
    └── tmp-outon4tl9z7
```

47 directories, 15 files

### 5 Scalability results

#### 5.1 On AWS

For the AWS experiments reported below, which were performed in 2019, we used the *then* most recent version of the runners: Cromwell 47 and Nextflow 19.04.1.5072

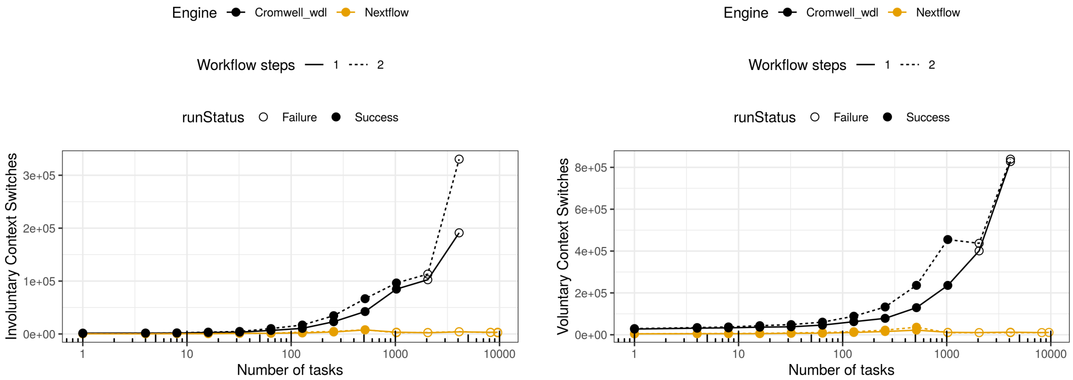

**Figure Supplementary 5: (Left) Involuntary and (Right) Voluntary context switches for each scalability scenario.**

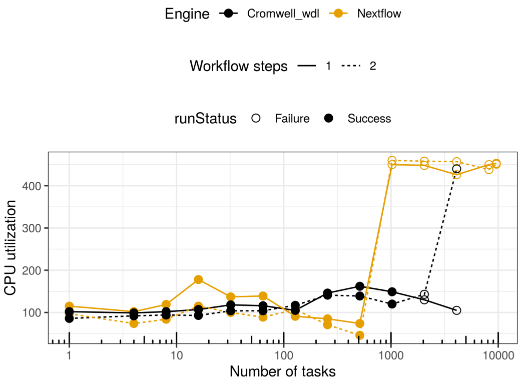

**Figure Supplementary 6: CPU utilization on the head node upon the execution of each workflow run.**

Cromwell was run with the in-memory database (the default), in run mode. Cromwell uses this database to track the execution of workflows and store outputs. For features like call caching, having a separate mysql database is necessary. This issue may have an effect on the CPU utilization.

#### 5.2 On Biocluster, Recent WfMS versions

The testing reported below was done in 2021, using the most recent version of the runners available: Cromwell 63, Toil 5.3.0 and Nextflow 21.04.1.5556. Experiments were performed on the `normal` queue of Biocluster, composed of 5 `Supermicro SYS-2049U-TR4` nodes, each of 72 cores. The cluster is not dedicated, so the data is affected by the queue load at the time.

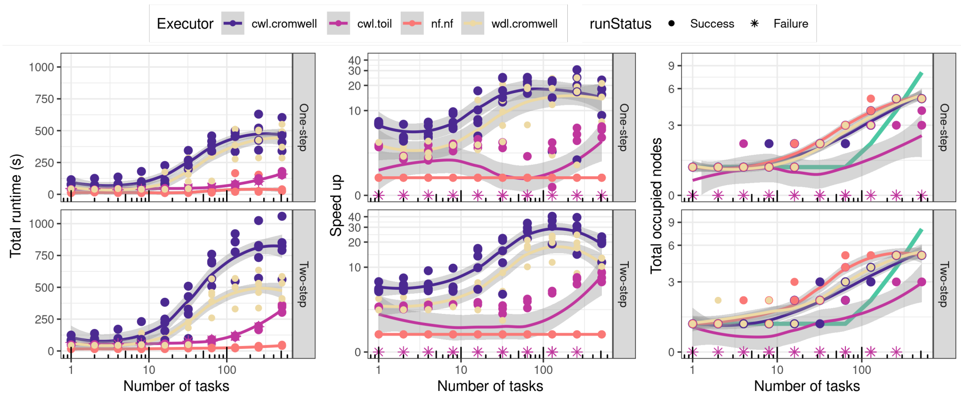

**Figure Supplementary 7: Scaling a one-step (top) and two-step (bottom) workflow in Toil+CWL, Cromwell+CWL, Cromwell+WDL and Nextflow. Nextflow can be up to 50x faster than Cromwell, regardless of the language (middle panel), while toil tends to fail unpredictably. The thick green line in rightmost panel is the theoretical number of cluster nodes, which is a ceiling of the ratio of the number of tasks divided by the number of cores per node (72)**

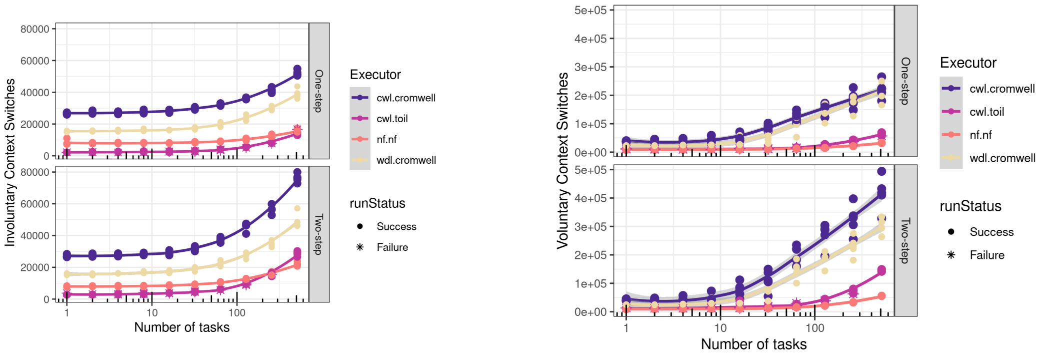

**Figure Supplementary 8: (Left) Involuntary and (Right) Voluntary context switches for each scalability scenario**

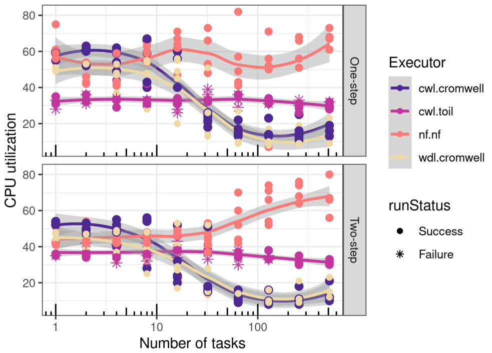

**Figure Supplementary 9: CPU utilization on the head node upon the execution of each workflow run**

#### 5.3 On Biocluster, Older WfMS versions

The experiments reported below were done in 2019, using the then most recent version of the runners: Cromwell 47 and Nextflow 19.04.1.5072. They were done on the `normal` queue, composed of 5 `Supermicro SYS-2049U-TR4` nodes, each of 72 cores. The cluster is not dedicated, so the data is affected by the queue load at the time.

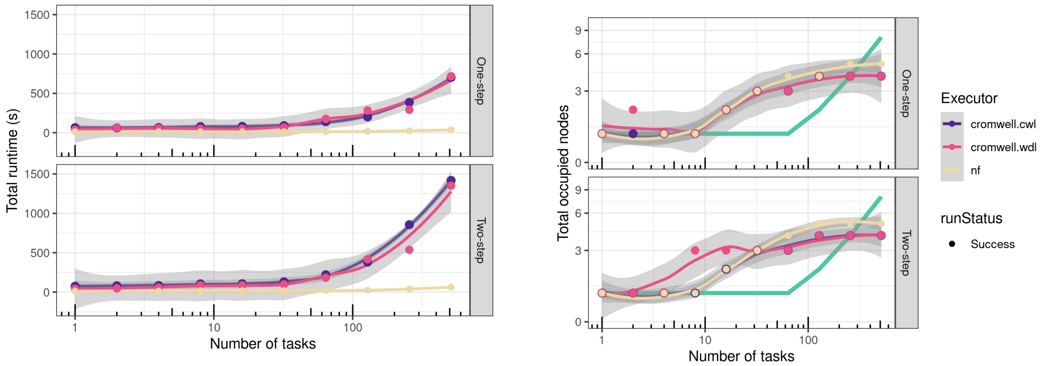

**(a) (Top) Execution times. (Bottom) Nextflow/Cromwell Speed-up.**

**(b) Tasks distribution across nodes.**

**Figure Supplementary 10: Scaling a one-step (solid line) and two-step (dashed line) workflow in Cromwell+CWL (black), Cromwell+WDL (yellow) and Nextflow (blue). Nextflow can be up to 20x faster than Cromwell, regardless of the language (Supplementary 10a, bottom). The thick green line in Supplementary 10b is the theoretical number of cluster nodes, which is a ceiling of the ratio of the number of tasks divided by the number of cores per node (72)**

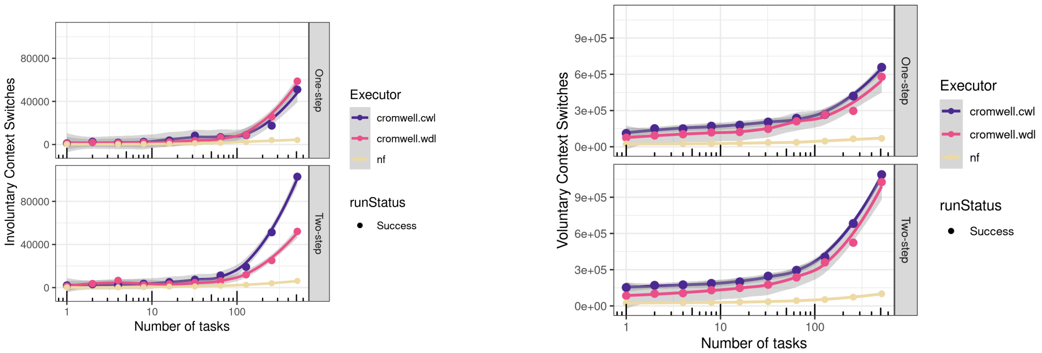

**Figure Supplementary 11: (Left) Involuntary and (Right) Voluntary context switches for each scalability scenario**

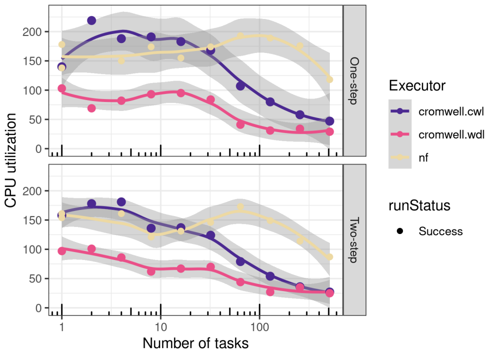

**Figure Supplementary 12: CPU utilization on the head node upon the execution of each workflow run**

#### Acronyms

**CWL** Common Workflow Languagen.

**DAG** Directed Acyclic Graph.

**GATK** Genome Analysis Toolkit.

**WDL** Workflow Description Language.

**WfMS** Workflow Management System.
